# Polyubiquitin-dependent recruitment of NEMO/IKKγ into T cell receptor signaling microclusters

**DOI:** 10.1101/617126

**Authors:** Elizabeth A. DeRiso, Andrea L. Szymczak-Workman, Angela Montecalvo, Joanne M. Murphy, Maria-Cristina Seminario, Lawrence P. Kane, Stephen C. Bunnell

## Abstract

The NF-κB essential modulator protein (NEMO) is required for activation of canonical NF-κB by the T cell antigen receptor (TCR). However, the subcellular localization of NEMO during this process is not well understood. By dynamically imaging fluorescent NEMO chimeras in live human T cells, we demonstrate that NEMO is rapidly recruited into TCR microclusters via domains previously implicated in the recognition of linear and K63-linked polyubiquitin. The recruitment of NEMO into TCR microclusters requires the activities of the tyrosine kinases Lck and ZAP-70, but not the adaptor proteins LAT or SLP-76. Thus, our findings reveal that the pathways leading from TCR to NF-κB bifurcate downstream of ZAP-70 to independently control the recruitment and activation of NEMO.

## Introduction

The non-catalytic adaptor subunit of the <u>i</u>nhibitor of <u>κ</u>B <u>k</u>inase (IKK) complex, <u>N</u>F-κB <u>e</u>ssential <u>mo</u>dulator protein - NEMO (or IKKγ) binds to the serine/threonine kinases IKKα and IKKβ (Israel, 2010). NEMO, like NF-κB itself, plays critical roles in T cell development and differentiation. Animal studies confirm that NEMO is required for T cell development beyond the double positive stage, the development of regulatory T cells, and the activation of canonical NF-κB isoforms downstream of the TCR. NEMO functions downstream of multiple receptors and is indispensable for the proper development and function of the vertebrate immune system (Hayden et al., 2006). In humans and mice, NEMO is X-linked, and null mutations are invariably lethal to males (Hanson et al., 2008; Orange et al., 2004). Males carrying hypomorphic alleles are unable to mount innate immune responses to bacterial pathogens, produce pathogen-specific antibodies, or generate effective T cell responses. Although female carriers appear normal, their peripheral lymphocytes exhibit fully non-random X chromosome inactivation, confirming that NEMO is essential for lymphocyte development.

NEMO is an alpha-helical dimer that contains segments known as the helix 1 (HLX1), coiled-coil 1 (CC1), helix 2 (HLX2), coiled-coil 2 (CC2), and leucine zipper (LZ) domains. NEMO interacts with IKKα and IKKβ via HLX1 (Napetschnig and Wu, 2013b). The CC2 and LZ domain comprise the <u>N</u>EMO <u>u</u>biquitin-<u>b</u>inding (NUB) domain, which binds linear polyubiquitin (pUb) chains (Laplantine et al., 2009; Lo et al., 2009; Rahighi et al., 2009). The helical region is linked to an unstructured linker and a C-terminal <u>z</u>inc <u>f</u>inger (ZF) domain. The NEMO ZF domain also binds pUb, but prefers K63-linked polymers (Cordier et al., 2009; Laplantine et al., 2009; Ngadjeua et al., 2013). Mutations that impair the interactions of NEMO with the IKK kinases, linear pUb chains, or K63-linked pUb chains are among the most common causes of NEMO-associated immune dysfunction (Hanson et al., 2008; Senegas et al., 2015).

NEMO mediates the activation of the IKK complex in response to the ubiquitination of several receptor-associated scaffolds. For example, Myd88-dependent Toll-like receptors (TLRs) and IL-1 family cytokine receptors employ <u>I</u>L-1 <u>r</u>eceptor <u>a</u>ssociated <u>k</u>inases (IRAKs), while TRIF-dependent TLRs and TNF family cytokine receptors employ <u>r</u>eceptor-<u>i</u>nteracting <u>p</u>rotein <u>k</u>inases (RIPKs). In each of these systems, an oligomeric scaffold is modified by ubiquitin E3 ligases to generate K63-linked and linear pUb chains (Tarantino et al., 2014). These chains mediate the recruitment of the NEMO-containing IKK complex and the <u>T</u>GFβ receptor-<u>a</u>ssociated kinase (TAK1), which phosphorylates IKKα and IKKβ (Napetschnig and Wu, 2013b). In contrast, lymphocyte antigen receptors activate the IKK complex via oligomeric ‘CBM’ complexes, which are composed of the CARD domain-containing adaptors <u>C</u>arma1 and <u>B</u>cl10, and the paracaspase <u>M</u>alt1.

While the localization of the IKK complex relative to IL-1 and TNF receptors has been addressed, the redistribution of the IKK complex in response to antigen receptor stimulation is poorly understood (Cheng et al., 2011; Tarantino et al., 2014). Early imaging studies indicated that NEMO enters the cSMAC within fifteen minutes (Rossman et al., 2006; Schaefer et al., 2004; Weil et al., 2003). Model systems in which T cell activation is achieved via surface-immobilized or lipid bilayer-tethered ligands offer enhanced spatial and temporal resolution. In both models, the TCR accumulates within discrete ‘microclusters’ that recruit and activate the TCR-proximal tyrosine kinase ZAP-70 (Bunnell et al., 2002; Campi et al., 2005; Yokosuka et al., 2005). In bilayer models, these TCR microclusters arise in the periphery of the nascent synapse, are transported to a CD28^hi^/TCR^lo^ signaling domain at the boundary of the cSMAC, and are eventually inactivated within the TCR^hi^ core of the cSMAC (Yokosuka et al., 2008). In contrast, when surface-immobilized ligands are employed, the proximal signaling complexes containing the TCR and ZAP-70 remain fixed in place (Bunnell et al., 2002). Proteins with critical roles in second messenger production and NF-ĸB activation (*e.g*. LAT, Gads, SLP-76, Vav1, and PLCγ1) are initially recruited to TCR microclusters, but segregate into distinct structures over time (Bunnell, 2010). These ‘SLP-76 microclusters’ are ultimately transported to the center of the contact where signaling is terminated (Hashimoto-Tane and Saito, 2016). The latter approach facilitates the allocation of signaling molecules into one of two signaling complexes: the TCR microcluster or the SLP-76 microcluster.

To date, no study has addressed the relationship of NEMO to these early forming signaling microclusters in response to TCR engagement (Cheng et al., 2011). Here, we report that immobilized TCR ligands trigger the recruitment of NEMO into structures that co-localize precisely with ZAP-70. Furthermore, our findings indicate that NEMO is recruited to non-degradative pUb chains assembled on the TCR, ZAP-70 or a tightly associated scaffold. We propose a revised model of TCR-dependent NF-κB activation in which the recruitment of NEMO to the TCR is a key event for optimal NF-κB activation.

## RESULTS

### NEMO is recruited into a pool of immobile microclusters in response to TCR ligation

To monitor the localization of NEMO within the immune synapse, we tagged human and murine NEMO with monomeric fluorescent proteins. When expressed at equivalent levels in Jurkat human leukemic T cells, these chimeras were both able to bind endogenous human IKKβ (**Supp. Figs. 1A-B**). The murine chimera also reconstituted PMA/ionomycin triggered NF-ĸB responses in the NEMO-deficient, Jurkat-derived, reporter line 8321 (**Supp. Fig. 1C**) (He and Ting, 2002). Because this reporter line expresses very low levels of the TCR, it could not be used to address receptor-triggered events.

To examine the dynamics of NEMO recruitment, Jurkat cells transduced with murine mYFP.NEMO chimeras were stimulated on functionalized glass substrates and imaged using spinning-disc confocal microscopy. Substrates coated with integrin ligands (*e.g*. fibronectin or recombinant human VCAM-1) induce T cell attachment and spreading. Under these conditions, NEMO was predominantly cytoplasmic (**Fig. 1A; Supp. Fig. 1E**). However, discrete structures of varying size were also present. These included: *i)* a perinuclear pool (cyan ‘P’), which was seen to undergo slow, continuous movements, *ii)* a pool of mobile objects of small to moderate size, and *iii)* NEMO macroclusters (red ‘M’), which moved erratically, but occasionally underwent rapid directional movements (Supplemental Movie 1). Substrates coated with TCR ligands (*e.g*. anti-CD3ε; OKT3) elicited all of the pools observed on integrin ligands alone, but also induced the formation of abundant, immobile NEMO microclusters, whether or not VCAM was present (**Fig. 1A; Supp. Movies 2-3; Supp. Fig. 1E**). Importantly, similar pools of NEMO were also observed in primary human T cell blasts transfected with vectors expressing an identical NEMO chimera (**Fig. 1B, Supp. Movie 4**).

**Figure 1:**
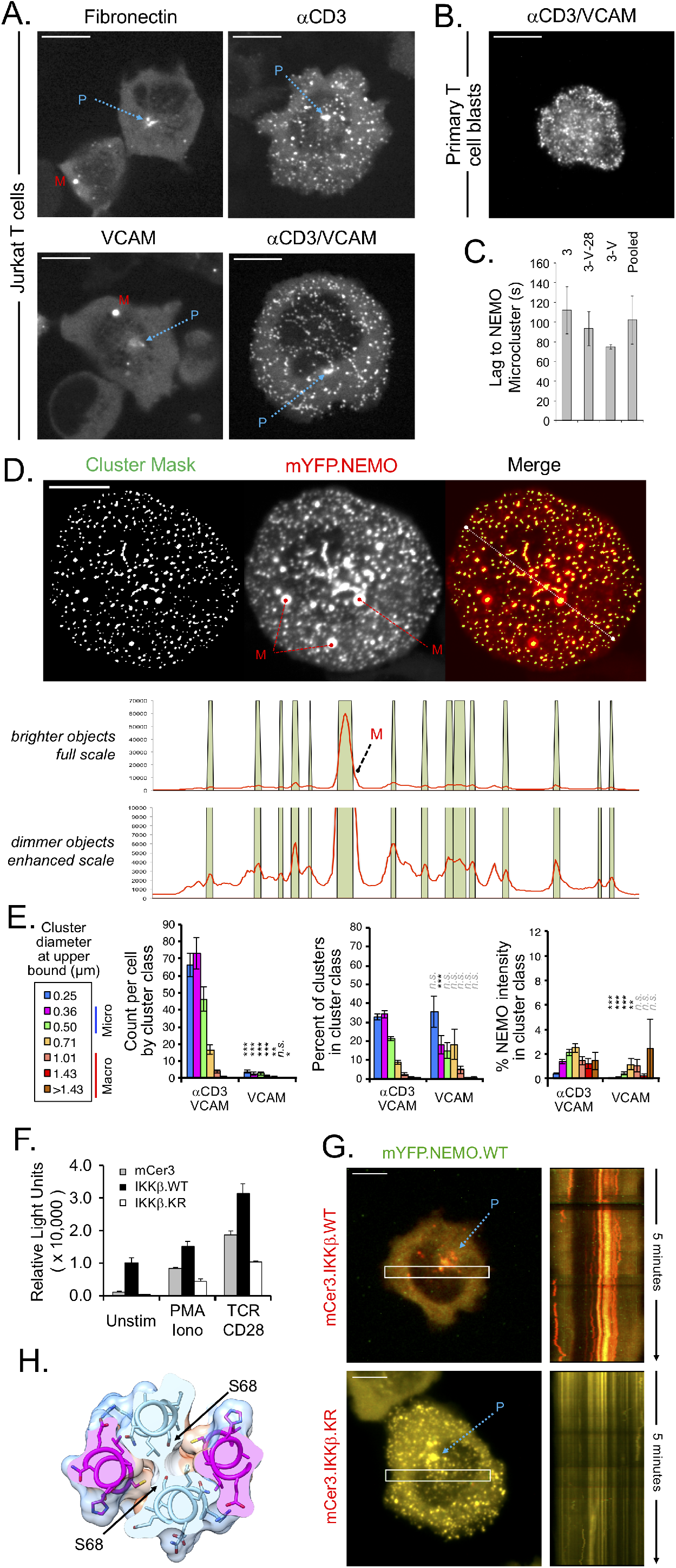
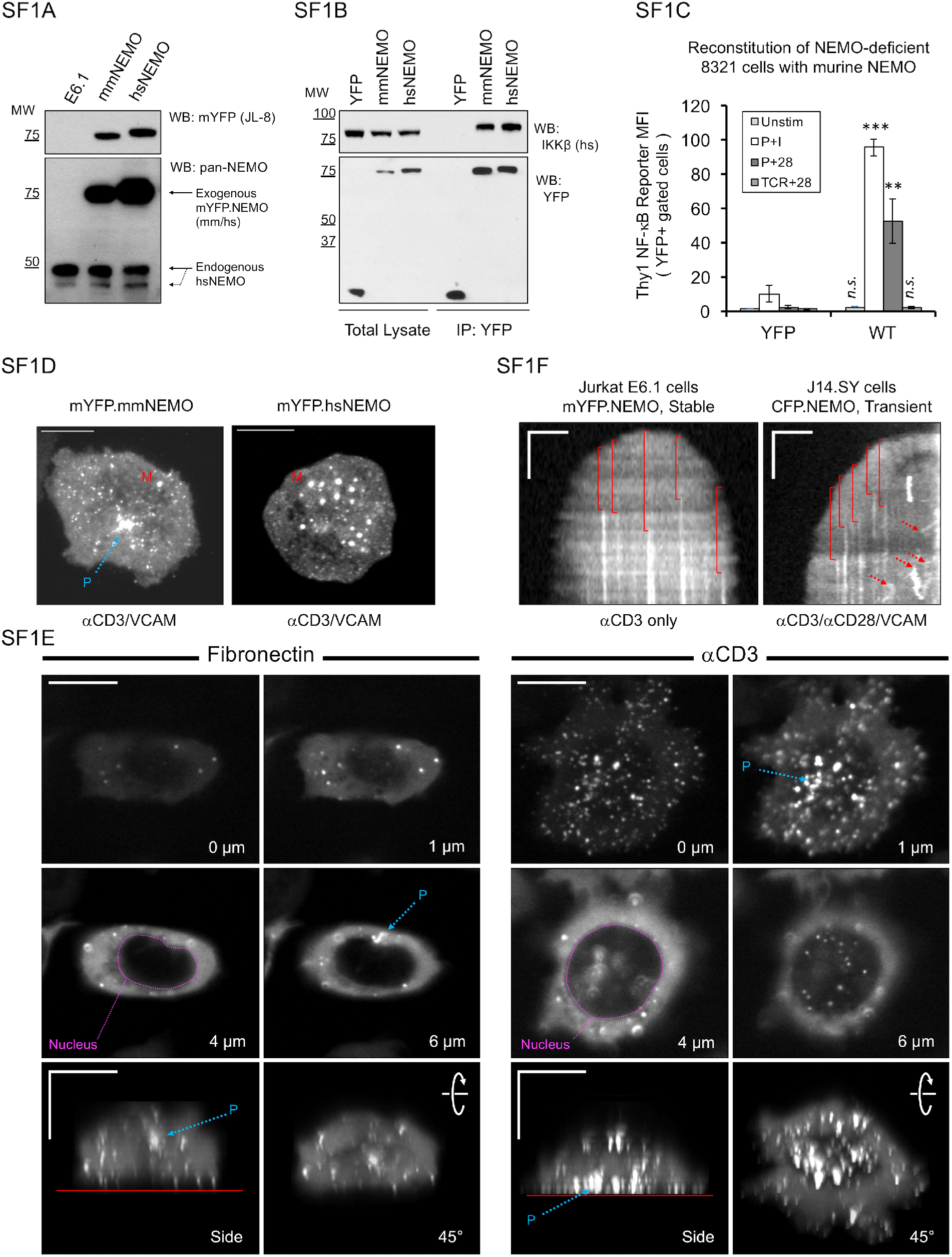
TCR stimulation triggers the rapid recruitment of NEMO into static structures associated with the stimulatory substrate. A) Jurkat T cells stably transduced with mYFP.NEMO were activated on stimulatory glass surfaces coated with 10 μg/ml anti-CD3 (OKT3), 1 μg/ml rhVCAM1 and/or 10 μg/ml fibronectin, as indicated. Dotted blue arrows, marked ‘P’, identify perinuclear pools of NEMO. Objects marked with a red ‘M’ are typical of the large objects referred to as ‘macroclusters’. Similar patterns were observed in every experiment (n≥3 for each condition). Scale bars: 10 μm. B) Primary human T cell blasts were transduced, stimulated, and imaged as in A. Images are representative of four experiments. C) The lag between local contact initiation and NEMO microcluster formation was calculated for cells captured in the process of spreading. Data are presented as the mean ± SD based on the number of cells examined. 468 clusters were observed in thirteen cells stimulated on anti-CD3 (3, n=8 cells) or anti-CD3 and rhVCAM1 (3-V, n=2 cells). No significant differences were observed between any group of cells and the pool of all conditions, or the cells stimulated on anti-CD3. D) Jurkat T cells stably transduced with mYFP.NEMO were stimulated on anti-CD3 and rhVCAM1 and fixed after five minutes. NEMO clusters were identified algorithmically (see Materials and Methods). The resulting masks are pseudo-colored (green) and superimposed on the raw image (red). Line scans show NEMO intensity (red line) along the indicated white line relative to the cluster masks (shaded green). E) Jurkat T cells were transiently transfected or stably transduced with mYFP.NEMO and imaged on the indicated substrates. NEMO clusters identified as in D were binned into classes based on cluster area. Classes are labeled using the diameter of a circle with an area equivalent to the upper bound of the class. Graphs present the cluster count (upper), the fractional distribution of clusters by class (middle), and the fraction of the NEMO intensity in the imaging plane that is captured within each class (lower). Data are presented as the mean ± SEM, based on the number of cells analyzed. Cluster quantitation was performed using 23 cells stimulated on anti-CD3 and rhVCAM1 and 13 cells stimulated on rhVCAM1 alone. Statistical differences among classes were determined using Student’s T-test: p < 0.05, *; p < 0.01, **; p < 0.001, ***. F) Jurkat cells expressing mYFP.NEMO and NFκB-luciferase plus either mCer3 itself or mCer3 chimeras with WT or kinase-deficient (KR) IKKβ were stimulated as indicated; levels of luciferase were measured to indicate the amount of NF-κB transcription. G) Jurkat T cells stably expressing mYFP.NEMO WT and either mCer3.IKKβ WT or mCer3.IKKβ-K44R (kinase-deficient mutant). Kymographs for each were taken from the region in the white box, and represent 5 minutes in time. H) Model showing the location of NEMO S68 in the context of a NEMO dimer; phosphorylation of this site is predicted to cause destabilization of the IKK complex and dissolution of the NEMO/ IKKβ microcluster.

When Jurkat cells expressing either the human or murine chimeras were stimulated via the TCR and VLA-4, we observed NEMO in discrete structures that varied considerably in both size and fluorescent intensity (**Supp. Fig. 1D**). The most prevalent structures were similar in size to TCR/ZAP-70 and LAT/SLP-76 microclusters (250-500 nm diameter), so we refer to these structures as NEMO microclusters (Bunnell et al., 2002; Nguyen et al., 2008). Much larger objects (>750 nm diameter) were also commonly observed, and were typically an order of magnitude brighter than the NEMO microclusters on a ‘per pixel’ basis; we refer to these objects as NEMO ‘macroclusters’.

Given the unique, and unexpected, relationship between static NEMO microclusters and TCR ligation, we examined the properties of these structures in greater detail. By observing cells caught in the process of contact expansion, we were able to determine the lag between local contact initiation and the formation of NEMO microclusters. As revealed by kymographs depicting the behavior of NEMO over time, these structures were typically formed *de novo*, apparently through the recruitment of NEMO from the cytoplasmic pool (**Supp. Fig.1F**). NEMO microclusters formed within 102 ± 24 seconds (mean ± SD) of contact with the stimulatory substrate, regardless of the presence or absence of costimulatory ligands (**Fig. 1C**).

To further quantify the different pools of NEMO that we observed, we developed a segmentation algorithm capable of identifying both NEMO microclusters and macroclusters (**Fig. 1D**). NEMO structures were then grouped by area and quantified. The upper bound of each group is presented using the effective diameter of a circle of equivalent area (**Fig. 1E**). The TCR-dependent induction of NEMO microclusters is evident in both the raw cluster count and the fraction of total NEMO intensity that is captured by each class of clusters (**Fig. 1E**; upper and lower panels; magenta and green bars). However, the fraction of total NEMO intensity within macroclusters was less profoundly impacted, consistent with our observation that these structures pre-exist and are not induced by TCR ligation (**Fig. 1E**; lower panel; orange, red, and brown bars).

### Negative feedback between activation of IKKβ and the assembly of NEMO microclusters

NEMO is thought to activate of IKKα and IKKβ (which then go on to phosphorylate IκBα) by delivering these kinases to poly-ubiquitin bearing scaffolds where they encounter the upstream kinase TAK1 (Napetschnig and Wu, 2013a). To determine whether IKKβ could be activated within TCR/NEMO microclusters, we tagged IKKβ with the fluorescent protein mCerulean3 (mCer3) and examined whether this chimera co-localizes with NEMO in stably transduced Jurkat T cells. However, the ectopic expression of wild-type IKKβ triggers the spontaneous activation of NF-κB responses (**Fig. 1F**) and abolishes the recruitment of NEMO into microclusters (**Fig. 1G**). Since IKKβ can phosphorylate NEMO on serine and threonine residues that antagonize the activation of NF-κB, we postulated that IKKβ-mediated phosphorylation inhibits the scaffolding functions of NEMO (Prajapati and Gaynor, 2002). Consistent with this hypothesis, a kinase-deficient form of the mCer3.IKKβ chimera co-localizes perfectly with NEMO in microclusters and vesicles (**Fig. 1G**) (Woronicz, Gao, Cao, Rothe, & Goeddel, 1997; Zandi, Rothwarf, Delhase, Hayakawa, & Karin, 1997). Phosphorylation by IKKβ at S68 of NEMO has been reported to disrupt the formation of NEMO dimers and/or the association of IKKβ with NEMO (**Fig. 1H**) (Palkowitsch et al., 2008). Overall, these observations suggest that NEMO and IKKβ are recruited to TCR/ZAP-70 microclusters, activated *in situ*, and then released following the phosphorylation of NEMO by IKKβ.

### NEMO specifically co-localizes with TCR-induced ZAP-70 microclusters

As introduced above, early T cell receptor signaling is associated with formation of distinct microclusters containing either ZAP-70 or the adaptor protein SLP-76. When mCFP.NEMO was imaged in Jurkat cells expressing SLP-76.YFP, NEMO microclusters formed adjacent to, but did not coincide with, SLP-76 microclusters (**Fig. 2A**, left panel). Instead, a immobile pool of NEMO overlapped extensively with ZAP-70-positive TCR microclusters in both Jurkat T cells and primary human T cell blasts (**Fig. 2A**, center and right panels). By imaging cells expressing both NEMO and ZAP-70 chimeras in the process of spreading, we observed that ZAP-70 microclusters formed approximately one minute before NEMO microclusters appear (**Fig. 2B** and **Supplemental movie 5**). In most cases, NEMO gradually accumulated within ZAP-70 microclusters, consistent with recruitment of NEMO from the cytoplasm (**Fig. 2B**, kymographs 1-2). The perinuclear NEMO-containing structures (cyan ‘P’) moved slowly and did not interact with NEMO microclusters. However, some mobile NEMO clusters appeared to arrest in the vicinity of static ZAP-70 microclusters (**Fig. 2B**, kymographs 1-3, asterisks). These NEMO clusters then typically departed after a brief pause (**Fig. 2B**, kymograph 3, white arrows). In some cases, a small amount of NEMO remained associated with the TCR/ZAP-70 microcluster after these stopping events (**Fig. 2B**, kymograph 3, fourth asterisk from top). Thus, our data suggest that mobile NEMO clusters may exchange material with static TCR/NEMO microclusters.

**Figure 2:**
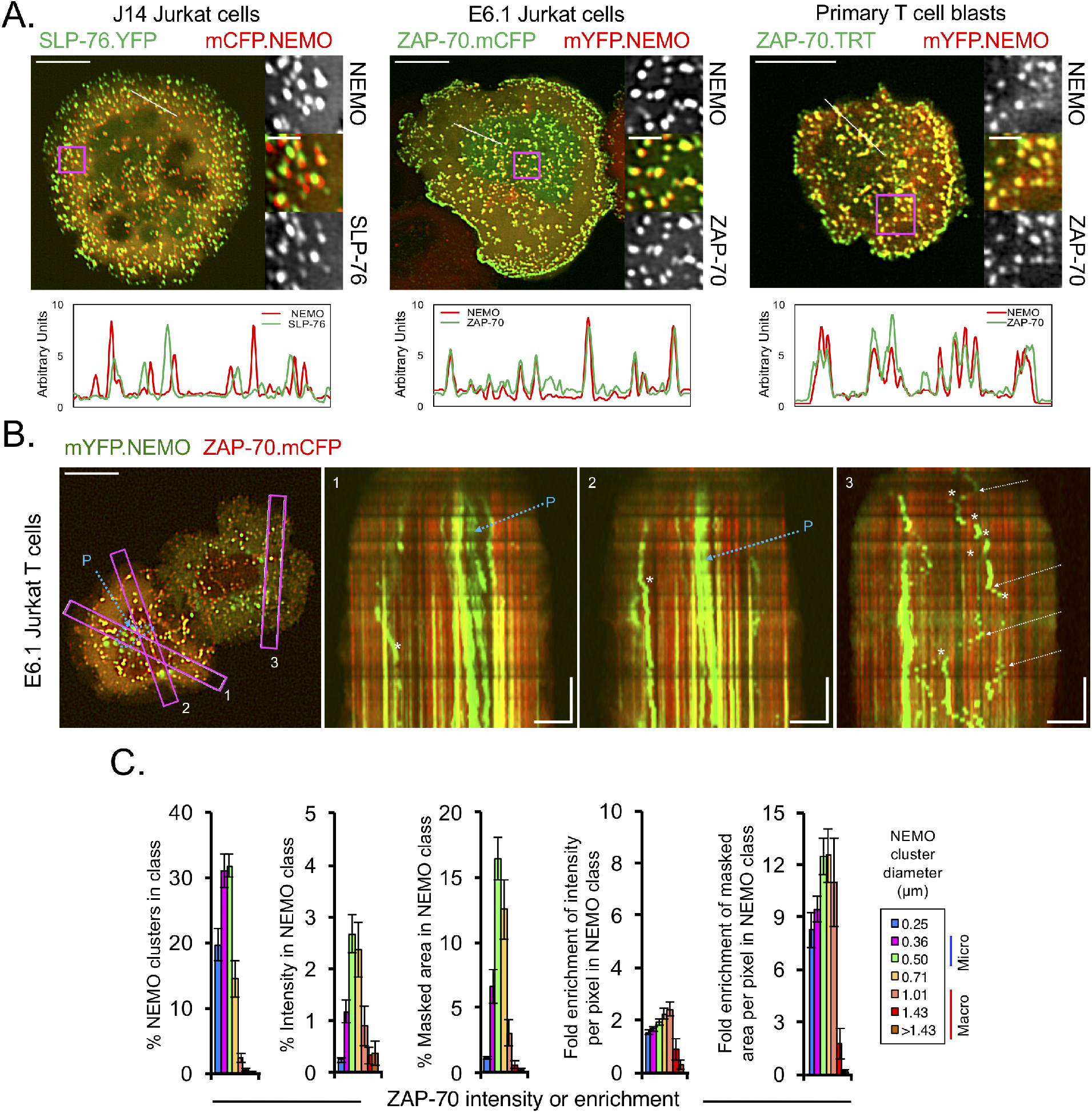
Static NEMO microclusters are preferentially associated with ZAP-70. A) SLP76-deficient J14 Jurkat cells stably reconstituted with SLP-76.YFP, parental Jurkat T cells and primary human T cell blasts were transiently transfected as indicated and stimulated on anti-CD3, rhVCAM1, and anti-CD28, as in Figure 1A. Images were pseudo-colored as indicated. The regions enclosed in magenta boxes are enlarged at right. Relative fluorescence intensities along the white diagonal lines are shown in the lower panels. Data are representative of three-six experiments for Jurkat cells and two experiments for primary cells. B) Jurkat T cells were transiently transfected with mYFP.NEMO and ZAP-70.mCFP, stimulated on anti-CD3, and imaged over time. A pseudo-colored still image is shown at left. Blue arrows identify a perinuclear pool that is not anchored to the substrate. The regions enclosed in magenta boxes were used to generate the kymographs shown at right. White arrows identify points at which static NEMO clusters begin to move, and asterisks identify the points at which mobile NEMO clusters stop. Still images are representative of six experiments. Scale bars: 10 μm (stills); 2 μm (insets); 5 μm × 60 seconds (kymographs). C) Graphs present the fractional distribution of NEMO clusters by size class (left), the fraction of total ZAP intensity within each NEMO class (center-left), the fraction of the total area masked as a ZAP cluster that lies within each NEMO class (center), the per-pixel enrichment of ZAP intensity within each NEMO class, relative to the entire cell (center-right), and the per-pixel enrichment of ZAP masked area within each NEMO class, relative to the entire cell (right). Statistical differences among corresponding classes were determined using Student’s T-test: p < 0.05, *; p < 0.01, **; p < 0.001, ***. Data are presented as the mean ±SEM, based on the number of cells analyzed. Calculations were performed for 12 cells co-expressing NEMO and ZAP-70.

To quantitate the degree of overlap of the various pools of NEMO with ZAP-70, we made further use of the masking algorithm described above. First, we determined the fraction of ZAP-70 intensity within each class of NEMO clusters, considering only the imaging plane (**Fig. 2C**, second panel from left). We performed similar calculations for the area masked as positive for ZAP-70 (**Fig. 2C**, third panel form left). Finally, we converted these values to enrichments per pixel, relative to the entire imaging field (**Fig. 2C**, rightmost panels). These data confirm that the smaller NEMO-containing structures preferentially overlap with ZAP-70, relative to the minimal overlap seen with the larger NEMO clusters.

### Structural requirements for NEMO localization in T cells

To determine which regions of NEMO control its localization in resting and activated T cells, we generated NEMO truncation mutants lacking the N-terminus, C-terminus, or both termini, with the latter construct containing only the “NUB,” or NEMO ubiquitin-binding domain (**Fig. 3A**). In functional assays, the N- and C-terminal deletion mutants both failed to reconstitute NF-κB activation in the NEMO-deficient reporter line (**Fig. 3B**). In cells plated on recombinant VCAM1 alone, the truncated chimeras were predominantly cytoplasmic, with few NEMO vesicles or macroclusters (**Fig. 3C**, top row). In response to TCR ligation, the truncated chimeras did not redistribute into the microclusters, mobile vesicles or macroclusters observed with wild-type NEMO (**Fig. 3C**, bottom row). A subset of cells expressing the NEMO-ΔC mutant formed macroclusters without developing microclusters, as reflected in our quantitative analyses (**Fig. 3D**, right panel). While perinuclear NEMO persisted in the NEMO-ΔC mutant, this pool was substantially reduced with the NEMO-ΔN and NEMO-NUB mutants.

**Figure 3:**
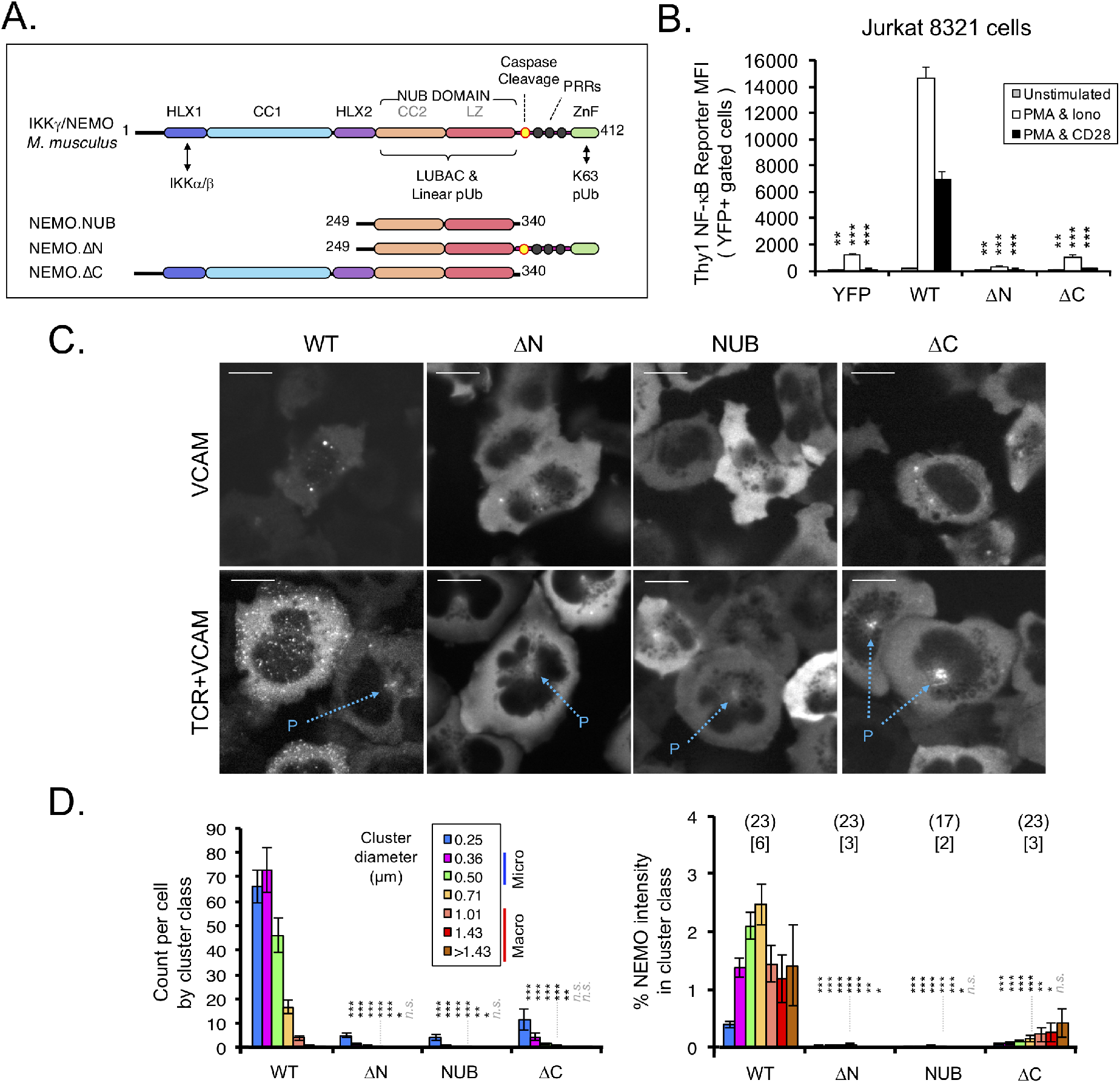
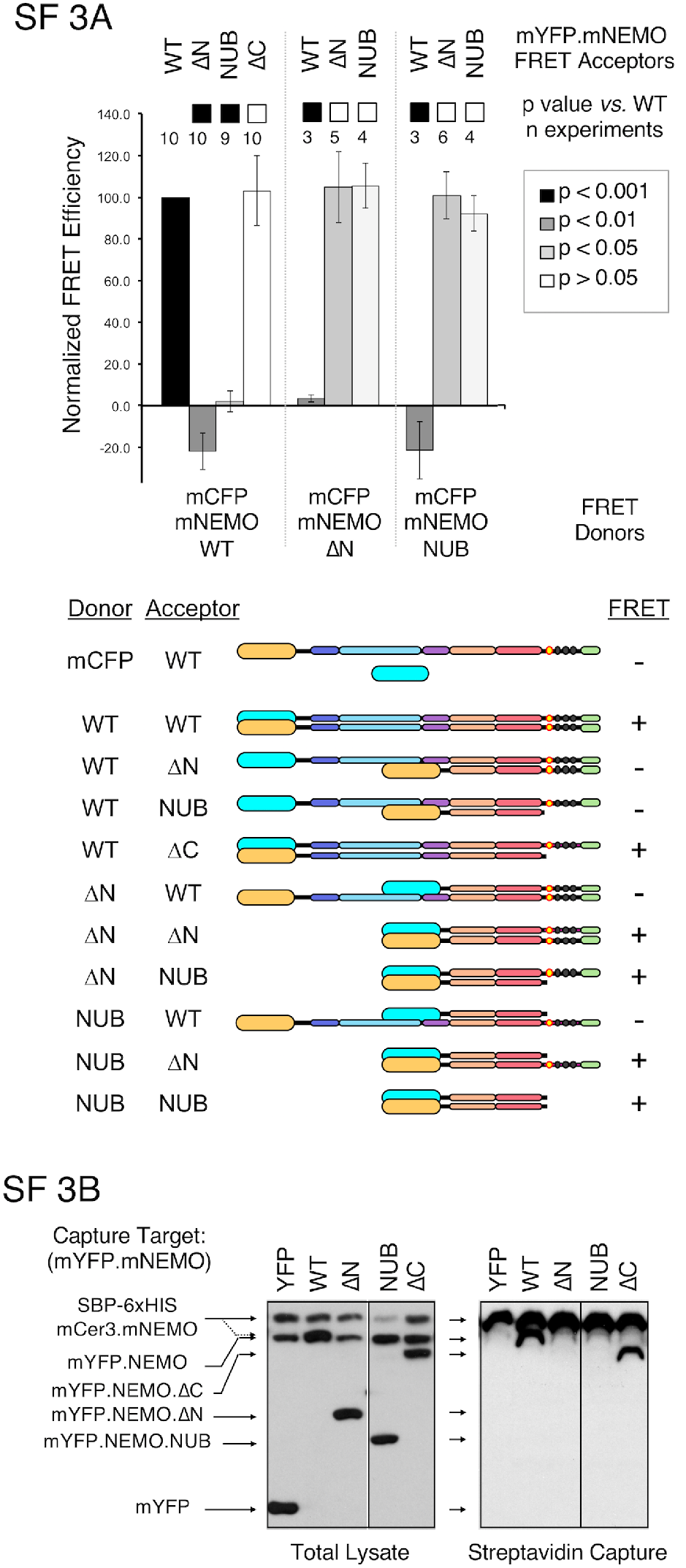
C-terminal truncations eliminate microcluster formation without impairing NEMO homo-dimerization. A) Diagram of NEMO domain structure and truncations used in subsequent experiments. B) NEMO-deficient 8321 cells expressing an NF-κB-driven rat Thy1 reporter were transiently transfected with vectors expressing mYFP alone, or mYFP.NEMO chimeras bearing the indicated deletions. After stimulation with PMA, plus either ionomycin or anti-CD28, these cells were analyzed by flow cytometry. Thy1 MFI is shown for cells expressing comparable levels of mYFP. Statistical differences were determined using Student’s T-test: p < 0.01, **; p < 0.001, ***. Data are presented as the mean ± SD of three technical replicates and are representative of 2-5 experiments. C) Jurkat T cells stably expressing wild-type or truncated mYFP.NEMO chimeras were stimulated as in Figure 1A and fixed after five minutes. Still images are representative of the cells quantitated in panel D. Scale bars: 10 μm. D) Transiently transfected and stably transduced Jurkat T cells expressing the indicated constructs were stimulated on anti-CD3 and rhVCAM1 and imaged live or after fixation. NEMO clusters were analyzed as in Fig. 1E. Data are presented as the mean ± SEM, based on the number of cells analyzed. Numbers in parentheses indicate the number of cells analyzed per condition; numbers in brackets indicate the number of independent experiments from which these cells were derived. Statistical differences among corresponding classes were determined using Student’s T-test: p < 0.05, *; p < 0.01, **; p < 0.001, ***.

Although fluorescence resonance energy transfer (FRET) experiments indicated that all three of the NEMO truncation variants form homotypic dimers in intact cells, the NEMO-NUB and NEMO-ΔN mutants failed to co-precipitate exogenous wild-type NEMO (**Supp. Figs. 3A-B**). This suggests that the NUB domain forms labile homodimers, and that the N-terminal helical domains are required to stabilize these dimers. Given the impact of the NEMO-ΔN and NEMO-NUB mutants on dimer stability, it is not surprising that these mutants fail to localize properly. However, the NEMO-ΔC mutant does not affect the stability of the NEMO dimer, and must impact the localization and function of NEMO via a distinct mechanism.

### Poly-ubiquitin binding sites promote assembly of NEMO into static microclusters

Given the known role of the NEMO ZF in pUb recognition, we generated NEMO chimeras bearing point mutations in the ubiquitin-binding sites of NEMO (**Fig. 4A**). The primary ubiquitin-binding site in the NUB domain recognizes both K63-linked and linear ubiquitin polymers, which is disrupted by a Y301S mutation (Ea et al., 2006; Lo et al., 2009; Rahighi et al., 2009; Yoshikawa et al., 2009). Binding of the NUB domain to linear ubiquitin is perturbed by the disease-causing R312Q mutant (Filipe-Santos et al., 2006; Hadian et al., 2011; Lo et al., 2009; Rahighi et al., 2009). The L322P mutation is within the leucine zipper region and prevents the recognition of both linear and K63-linked di-ubiquitin (Wu et al., 2006). Finally, the C-terminal ZF domain, which preferentially recognizes K63-linked ubiquitin polymers, is inactivated by the V407S mutant (Cordier et al., 2009; Laplantine et al., 2009; Ngadjeua et al., 2013). Thus, each of these mutants impaired the reconstitution of NFκB activation in a NEMO-deficient reporter cell line: the Y301S and L322P mutants were profoundly inactive, the R312Q mutant was impaired and the V407S mutant showed an intermediate defect (**Fig. 4B**). Based on FRET and co-immunoprecipitation assays, the L322P mutation disrupted NEMO dimerization and was not pursued further. None of the other mutants inhibited the assembly of NEMO dimers or the interaction of NEMO with IKKβ, strongly suggesting that they affect NEMO function via alternative mechanisms (Supp. Figs. **4A-C**). When Jurkat T cells expressing each of the remaining mutants were stimulated via the TCR, the static microclusters observed with wild-type NEMO were absent (**Fig. 4C; Supp. Movies 6-9**). However, macroclusters were more pronounced in the context of the Y301S and R312Q mutations and were somewhat reduced in the context of the V407S mutation (**Fig. 4D**). The wild-type and Y301S NEMO chimeras behaved similarly in transiently transfected primary human T cell blasts, with wild-type NEMO entering TCR/ZAP-70 microclusters, while the Y301S mutant remained in the cytoplasm and in pre-existing macroclusters (**Fig. 4E; Supp. Fig. 4D**).

**Figure 4:**
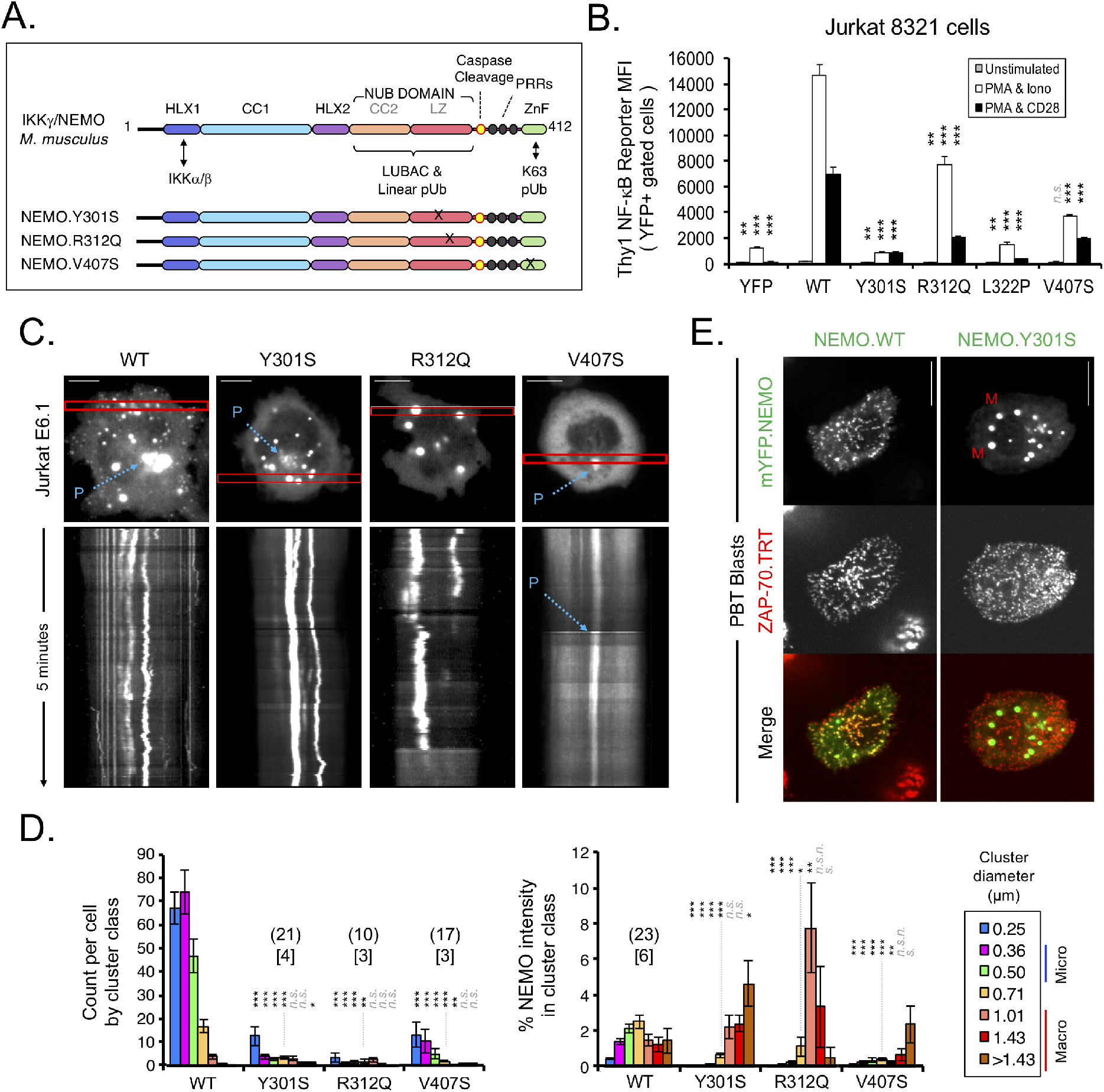
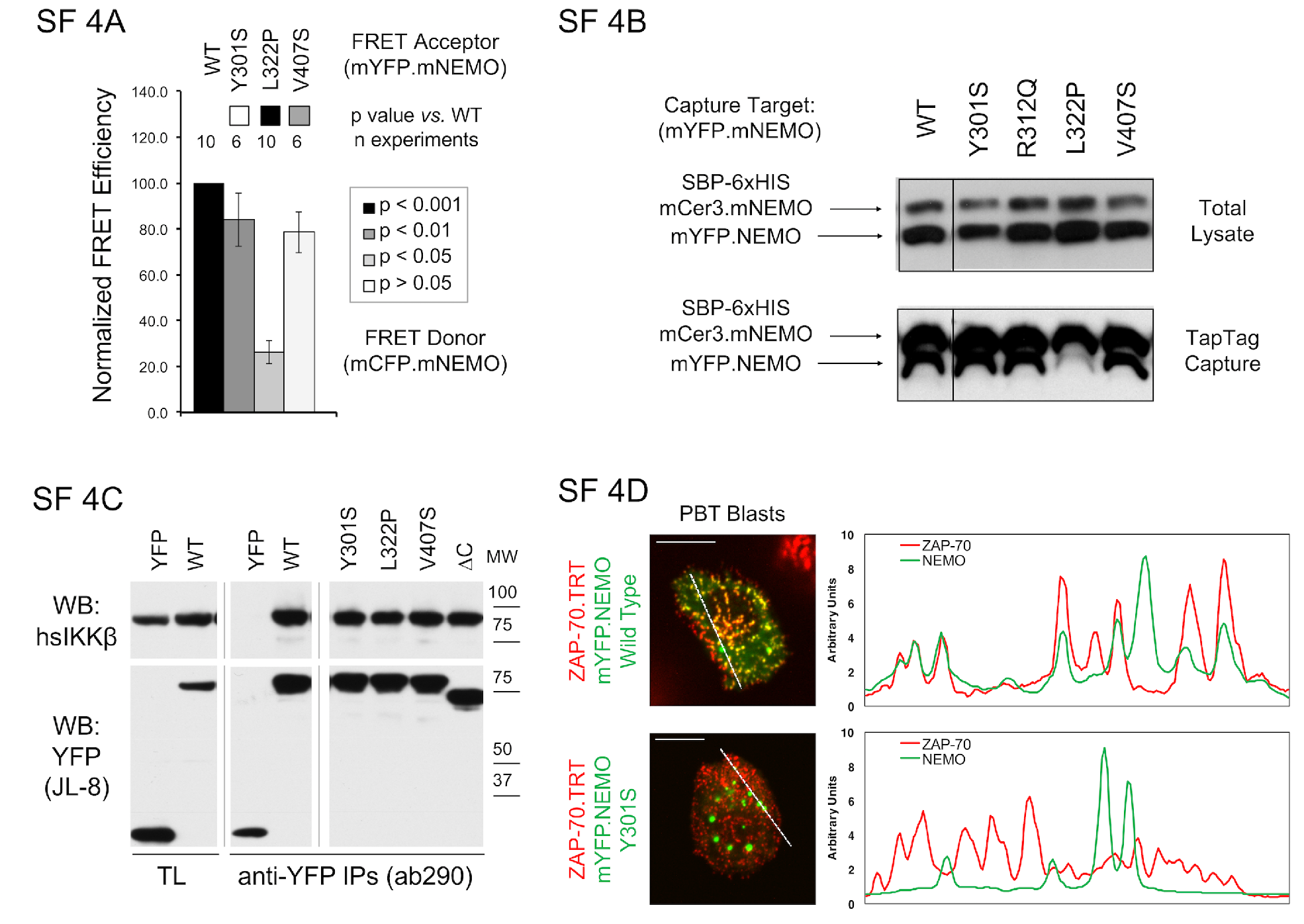
NEMO ubiquitin binding links NEMO microcluster recruitment to NF-κB activity. A) Diagram of NEMO point mutations used in subsequent experiments. B) The functionality of mYFP.NEMO chimeras bearing mutations that impair poly-ubiquitin binding was assessed as in Fig. 3. Data are presented as mean ± SD of 3 technical replicates and are representative of 3-5 experiments. C) Jurkat T cells stably expressing wild-type or mutant mYFP.NEMO chimeras were stimulated on anti-CD3 and rhVCAM1-coated substrates and imaged continuously for at least 5 minutes. Still images are representative of the cells quantitated in panel D. Kymographs were derived from the sub-regions boxed in red. Scale bars: 10 μm. D) Jurkat T cells expressing the indicated constructs were stimulated on anti-CD3 and rhVCAM1. NEMO clusters were imaged, analyzed, and presented as in Fig. 2. E) Primary human T cell blasts were transfected with vectors encoding either a wild-type or a Y301S mutant mYFP.NEMO chimera and a ZAP-70.TRT chimera. Transfected cells were stimulated on anti-CD3, rhVCAM1, and anti-CD28, fixed after 5 minutes, and imaged as above. Images are representative of 4 experiments for NEMO.WT and 2 experiments for NEMO.Y301S. Scale bars: 10 μm.

### NEMO microcluster formation requires the catalytic activities of Lck and ZAP-70

To clarify the molecular mechanisms underlying NEMO recruitment into TCR/ZAP-70 microclusters, we assessed the role of TCR-proximal signaling cascades, using both genetic and pharmacological approaches. Thus, Jurkat-derived T cell lines lacking the Src-family kinase Lck (J.CaM1.6) developed very few NEMO microclusters in response to TCR-stimulation, even though NEMO macroclusters were clearly present (**Fig. 5A**). Importantly, reconstitution with a fluorescently-tagged Lck chimera restored the formation of NEMO microclusters (**Fig. 5A**). The Src-family kinase inhibitor PP2 did not impair the formation of TCR microclusters, but is known to prevent the phosphorylation of the TCR and the recruitment of ZAP-70 (Bunnell et al., 2002; Hanke et al., 1996). In addition, PP2-treated cells formed fewer static NEMO microclusters in response to TCR ligation, even though NEMO vesicles and macroclusters were still present (**Figs. 5B-C; Supp. Movies 10-11**). A similar effect was observed with the ZAP-70/Syk inhibitor piceatannol (**Fig. 5C**, right – “Pic”). We also examined the localization of NEMO in the Jurkat-derived P116 cell line, which lacks endogenous ZAP-70 (Williams et al., 1998). When P116 cells expressing a wild-type ZAP-70 chimera were stimulated on immobilized TCR ligands, the NEMO chimera was recruited into TCR/ZAP-70 microclusters, mobile vesicles and macroclusters (**Fig. 5D**, left column; **Supp. Movie 12**). However, in P116 cells expressing a kinase-inactive variant of ZAP-70 (K369R), NEMO primarily entered macroclusters and was largely excluded from TCR/ZAP-70 microclusters (**Fig. 5D**, right column; **Supp. Movie 13**). These results confirm that the catalytic activity of ZAP-70 is required for the recruitment of NEMO into TCR microclusters.

**Figure 5:**
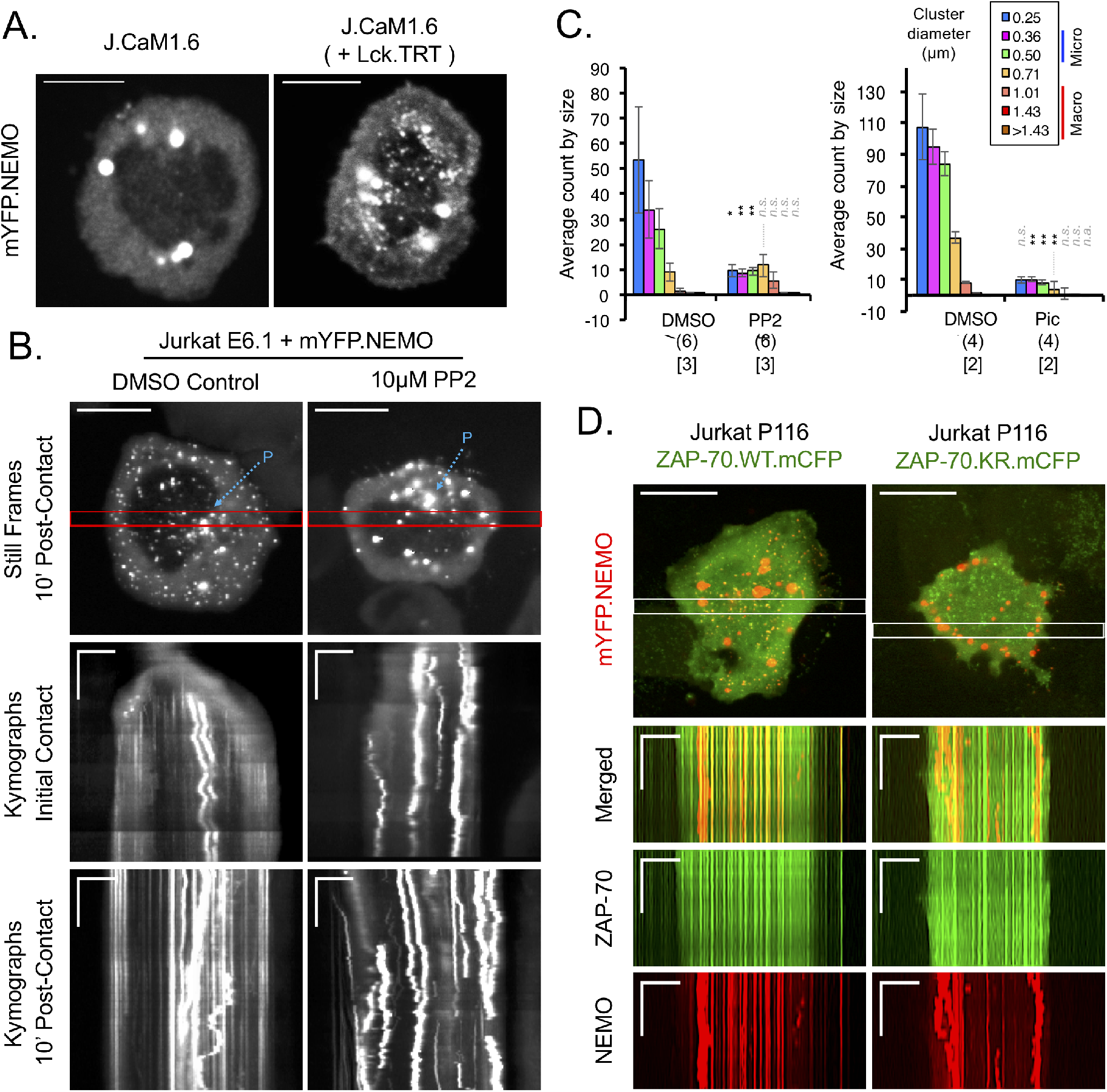
The catalytic activities of Lck and ZAP-70 are required for the assembly of NEMO microclusters. A) Lck-deficient J.CaM1.6 Jurkat cells were transiently transfected with vectors encoding either mYFP.NEMO alone (left) or mYFP.NEMO and wild-type Lck.Myc.TRT (right), stimulated on substrates coated with anti-CD3 and rhVCAM1, and fixed after 5 minutes. Still images are representative of the cells examined in panel B. B) Jurkat T cells stably expressing wild-type mYFP.NEMO were pre-incubated with 10μM PP2 or a corresponding volume of DMSO for 10 minutes. Cells were stimulated on surfaces coated with anti-CD3 and rhVCAM1 in the presence of the corresponding compounds. Images were acquired continuously for five minutes, beginning two and ten minutes after the addition of cells. Still images derived from the later series are representative of the cells quantitated in panel C. Kymographs derived from both acquisitions are shown. The later kymograph is derived from the region boxed in red. C) Jurkat T cells stably expressing wild-type mYFP.NEMO were pre-treated with 10μM PP2, 100μM piceatannol, or independent DMSO controls, and stimulated on anti-CD3 and rhVCAM1. NEMO clusters were imaged and analyzed as in Fig. 2. D) ZAP-70-deficient P116 Jurkat T cells were transiently transfected with vectors encoding mYFP.NEMO and either a wild-type (WT) or a kinase-dead (K369R, KR) ZAP-70.mCFP chimera. Cells were stimulated on substrates coated with anti-CD3, rhVCAM1, and anti-CD28 and imaged continuously for five minutes. Still images are shown above. Kymographs are derived from the regions boxed in white. Images are representative of two experiments. Scale bars: 10 μm (stills); 5 μm × 60 seconds (kymographs). See Figure 7B for NEMO cluster quantitation in J.CaM1.6 and P116 cells.

### NEMO microcluster recruitment does not require the LAT, SLP-76 or Carma1 adaptors

ZAP-70 activation promotes the generation of second messengers via signaling complexes nucleated by the adaptor proteins LAT and SLP-76 and activation of PLCγ1, leading to induction of transcription factors like NFAT, AP-1 and NF-κB. Therefore, we were surprised to find that Jurkat-derived cell lines lacking LAT (J.CaM2) or SLP-76 (J14) still developed static NEMO microclusters with the same size distribution as parental WT Jurkat T cells (**Figs. 6A-B**). One of the second messengers generated by PLCγ1, DAG, activates PKCθ (Dienz et al., 2003; Herndon et al., 2001). PKCθ then initiates the assembly of CBM complexes, which are thought to be the primary link between the TCR and the IKK complex (Hayden and Ghosh, 2012; Paul and Schaefer, 2013; Thome et al., 2010). Again, we observed that NEMO microclusters appeared to form normally in Jurkat-derived JPM50.6 cells, which lack expression of the Carma1 adaptor, a key component of the CBM complex (**Figs. 6A-B**). Quantitative analyses of fixed cells verified that Lck and ZAP-70 are indeed crucial for the formation of NEMO microclusters, despite the lack of a requirement for the above adaptor proteins (**Fig. 6B**). To validate findings obtained with the JPM506 cells, we co-expressed and imaged NEMO and ZAP-70 chimeras in live JPM50.6 cells. These studies confirmed that the NEMO clusters were associated with static TCR/ZAP-70 microclusters (**Fig. 6C**, left), and that these structures were fixed in-place, as shown by kymograph analysis (**Fig. 6C**, right).

**Figure 6:**
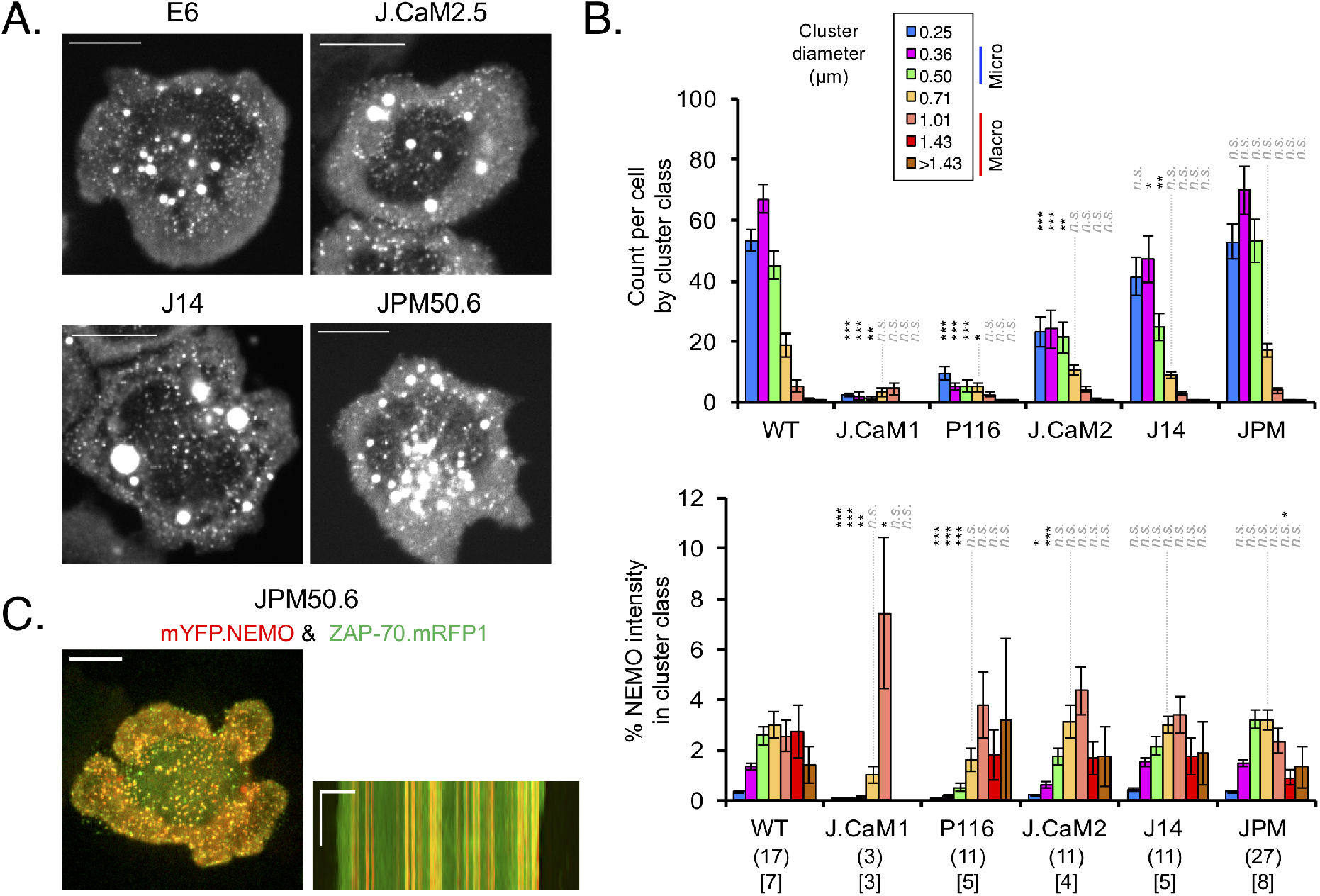
The formation of TCR-associated NEMO microclusters does not require the adaptors Carma1 or SLP-76. A) Wild-type Jurkat T cells and mutant lines lacking LAT (J.CaM2.5), SLP-76 (J14) or Carma1 (JPM50.6) were stably transduced with wild-type mYFP.NEMO, stimulated on anti-CD3 and rhVCAM1, and fixed after five minutes. Still images are representative of the cells examined in panel B. B) Mutant cell lines were prepared and stimulated as in A. NEMO clusters were imaged and analyzed as in Fig. 2. C-D) JPM50.6 Jurkat T cells were transiently transfected with vectors expressing fluorescent chimeras of NEMO and Zap70, and then stimulated, fixed, and imaged as in A. Representative still images and kymographs are shown (n=9 cells in 2 experiments). C) JPM50.6 cells were transfected with vectors expressing mYFP.NEMO and Zap70.mRFP1. Images were acquired continuously for five minutes, beginning two or 20 minutes after the addition of cells. Still images acquired early in each series are shown at left, with kymographs derived from the boxed regions at right (n=4 cells in 2 experiments for each condition). Scale bars: 10 μm (stills); 5 μm × 60 seconds (kymographs).

## Discussion

Although the roles of NEMO in T cell development and function have been extensively studied, the biochemical and cell-biological mechanisms that control the initial recruitment of NEMO and the activation of the IKK complex in response to TCR ligation have not been fully resolved (Cheng et al., 2011; Shi and Sun, 2015; Thome et al., 2010). While NEMO clearly enters antigen-induced immunological synapses, NEMO has never been visualized in real time during initial T cell contact formation (Hara et al., 2004; Shambharkar et al., 2007; Weil et al., 2003). This is important, because early signaling events downstream of the TCR precede formation of the immune synapse (Lee et al., 2002). Although NEMO enters multiple sub-cellular pools, our studies indicate that the ability of NEMO to activate NF-ĸB is specifically correlated with the ability of NEMO to enter TCR microclusters. The rapid recruitment of NEMO into these structures requires the pUb binding properties of NEMO and the activity of the TCR-associated tyrosine kinases Lck and ZAP-70. While IKK activation is typically placed downstream of the CBM complex, proximal signaling adaptors and those required for the assembly of CBM complexes, including LAT, SLP-76, and Carma1 are dispensable for the recruitment of NEMO into TCR microclusters. Thus, we propose that IKK complexes enter microclusters via TCR-proximal and CBM-independent linear and K63-linked pUb scaffolds that are assembled in response the activation of ZAP-70 (see model in **Fig. 7**).

**Figure 7:**
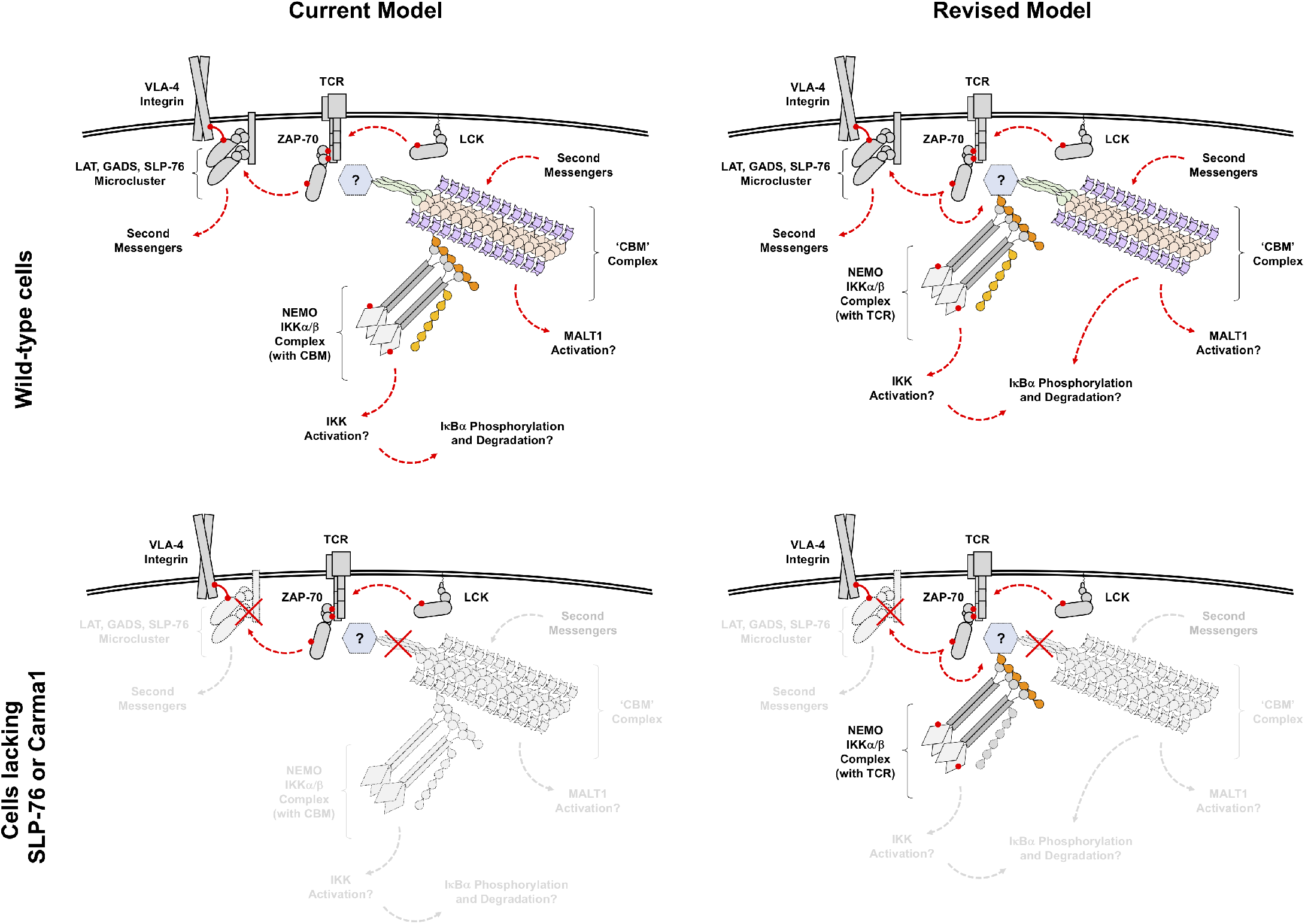
A revised model of TCR-dependent IKK activation. Conventional (left) and revised (right) models of IKK complex recruitment and activation downstream of the TCR. Conventional models place the polyubiquitin scaffolds to which NEMO is recruited on the CBM complex; the phosphorylation of IKKα and IKKβ and the polyubiquitination of NEMO are presumed to occur at this site (top left). The old model is incompatible with the observation that NEMO microclusters are unaffected by mutations that preclude the assembly of CBM complexes (bottom left). Our revised model places the polyubiquitin scaffolds to which NEMO is recruited on an as yet undefined component of the TCR/ZAP-70 microcluster (top right). This provides a potential explanation for the Carma1-independence of NEMO microclusters and for the retention of IKKβ phosphorylation in the absence of Carma1. However, this model still places Carma1 upstream of NEMO ubiquitination and I<u>κ</u>B degradation (bottom left).

NEMO protein is found in several intracellular pools in addition to microclusters. Large, membrane-bounded NEMO ‘macroclusters’ are relatively rare, but are very bright, and contain a significant fraction of cellular NEMO. These structures are present in unstimulated T cells, and are more abundant in the context of the Y301S and R312Q mutations, which hinder TCR-induced NF-ĸB activation and interfere with the ability of the NUB domain to bind linear pUb chains (Ea et al., 2006; Lo et al., 2009; Rahighi et al., 2009; Wu et al., 2006). Consequently, these macroclusters are unlikely to play a positive role in T cell activation. Because these structures fuse with one another and do not shed NEMO-containing structures, they may correspond to a degradative compartment, such as a lysosome. NEMO is also present in smaller clusters that are capable of moving within the cytoplasm. A subset of these structures moves in a manner consistent with the microtubule organizing center. We have not observed this population of perinuclear clusters interacting with any other pool of NEMO. Finally, many NEMO clusters are capable of rapid, directed movements through the cytoplasm, interacting with NEMO microclusters and occasionally fusing with membrane-bounded macroclusters. These observations indicate that NEMO enters post-endocytic compartments and suggest that NEMO may influence endocytic processes.

The IKK complex is a stable tetrameric structure that typically contains one NEMO homodimer and one dimer containing IKKα and/or IKKβ (Drew et al., 2007; Solt et al., 2009; Solt et al., 2007). Crystallographic data confirm that the HLX1, HLX2, and NUB domains all form dimers (Bagneris et al., 2008; Fujita et al., 2014; Grubisha et al., 2010; Rahighi et al., 2009; Rushe et al., 2008; Yoshikawa et al., 2009). Due to their atypical helical compositions and low melting temperatures, these isolated fragments of NEMO form weak dimers that are stabilized by flanking coiled regions or by ligand binding (Drew et al., 2007; Grubisha et al., 2010). Here, we have confirmed that wild-type and N-terminally truncated NEMO mutants form dimers in intact cells, but that these dimers cannot persist in the absence of the N-terminal helical region. The truncations that destabilize the NEMO dimer abolish most NEMO structures, including the perinuclear pool, which is otherwise unaffected by mutations impacting either the C-terminus or ubiquitin-binding capacity of NEMO. In contrast, dimer disruption cannot account for loss of microcluster formation by the ΔC, Y301S, R312Q, and V407S mutants, which can be coprecipitated with wild-type NEMO.

Our functional and imaging studies indicate that the ability of NEMO to enter microclusters in response to pUb recognition plays a crucial role in TCR-induced NF-κB responses. The NUB and ZF domains of NEMO have been reported to respectively prefer linear and K63-linked polymers (Hadian et al., 2011; Kensche et al., 2012; Laplantine et al., 2009; Lo et al., 2009; Ngadjeua et al., 2013). Mutation of each of these domains impacts the localization of NEMO in a distinct manner. Thus, mutations that inhibit linear pUb recognition by the NUB domain shift NEMO from microclusters into macroclusters, while those that eliminate or disable the ZF disrupt microclusters, macroclusters and vesicles, leaving only cytoplasmic and perinuclear pools of NEMO. Therefore, we propose that ZF-dependent K63-linked pUb recognition recruits NEMO to TCR-associated scaffolds, but predisposes NEMO to form macroclusters. In contrast, recognition of linear pUb by the NUB may promote sustained activation of IKKα/β by hindering the recruitment of NEMO into macroclusters.

Antigen-stimulated T cells form Carma1-dependent oligomeric signaling complexes that contain Bcl10 and Malt1. These CBM-containing structures, which are also known as <u>p</u>unctate and <u>o</u>ligomeric <u>k</u>illing or <u>a</u>ctivating <u>d</u>omains <u>t</u>ransducing <u>s</u>ignals (POLKADOTS), typically appear 10-20 minutes after synapse formation and are often found at sites distal to the contact interface (Paul et al., 2012; Paul et al., 2014; Rossman et al., 2006; Schaefer et al., 2004). Although these complexes are thought to link the TCR and IKK complex, these polymeric structures develop too slowly to account for the rapid recruitment of NEMO into the immunological synapse or initial activation of the IKK complex (Coornaert et al., 2008; Hara et al., 2004; Rebeaud et al., 2008; Ruland et al., 2003).

Activation of the IKK complex is associated with modification of NEMO by K63-linked and linear pUb chains (Deng et al., 2000; Fujita et al., 2014; Sun et al., 2004; Tokunaga et al., 2009; Zhou et al., 2004). However, phosphorylation of IĸBα by the IKK complex requires distinct inputs that independently control NEMO polyubiquitination and IKKα/β phosphorylation (Cheng et al., 2011). In particular, TCR-dependent polyubiquitination of NEMO is selectively eliminated in Carma1-deficient cells, while phosphorylation of IKKα/β by TAK1 can be perturbed without affecting NEMO polyubiquitination (Shambharkar et al., 2007; Srivastava et al., 2010). These observations are at odds with the view that the CBM complex is the sole scaffold responsible for the activation of the IKK complex. Our findings suggest that TCR microclusters can function as Carma1-independent platforms for NEMO recruitment. Since NEMO microcluster formation relies on the ZF domain, K63-linked ubiquitin scaffolds are likely to be produced within TCR microclusters. This would enable the simultaneous recruitment of TAK1 and NEMO to TCR-associated K63-pUb scaffolds and would provide a viable explanation for Carma1-independent IKKα/β phosphorylation.

Our observations suggest an updated model of antigen receptor-dependent NF-κB activation that parallels the events downstream of the receptors for IL-1β and TNFα. In all three systems, NEMO is recruited into receptor-associated micro-structures that form within minutes of receptor engagement (Tarantino et al., 2014). Photobleaching studies indicate that the cytokine-induced NEMO puncta are dynamic in composition, which is also true of the ZAP-70 present within TCR microclusters (Bunnell et al., 2002). Further, all three systems generate oligomeric assemblies of kinases that serve as substrates for the formation of ubiquitin polymers. Since chimeras that join the IKKα/β binding domain of NEMO to the tandem SH2 domains of ZAP-70 are sufficient to activate NF-ĸB following TCR ligation, the oligomeric nature of these scaffolds may contribute to the activation of the IKK complex (Weil et al., 2003). Finally, these receptor-associated NEMO puncta are reduced following the disruption of linear or K63-linked ubiquitin polymers or the disabling of the corresponding ubiquitin binding domains in NEMO. Based on these parallels, it is likely that TCR-proximal microcluster components serve as substrates for the formation of the polyubiquitin chains that recruit and activate the IKK complex.

## Materials and Methods

### Reagents

Recombinant proteins, biologicals, and antibodies were obtained as follows: recombinant human VCAM-1 (#862-VC) from R&D Systems (Minneapolis, MN); recombinant human IL-2 from the NIH AIDS Reagent Program; human plasma fibronectin (#2334564) and phytohemagglutinin (#10576015) from Invitrogen (Carlsbad, CA); anti-human CD3ε (OKT3, BE0001-2; RRID:AB_1107632) and CD28 (9.3, BE0248; RRID:AB_2687729) from Bio X Cell (Lebanon, NH); anti-human TCR Vβ8 (C305) from hybridoma supernatants; anti-*A. v*. GFP (JL-8; RRID:AB_10013427) from Takara Bio (Kusatsu, Shiga Prefecture, Japan); anti-GFP (ab-290; RRID:AB_303395) from Abcam (Cambridge, MA); anti-IKKβ (2C8, #2370; RRID:AB_2122154) from Cell Signaling Technology (Danvers, MA); anti-NEMO/IKKγ (FL-419, sc-8330; RRID:AB_2124846) from Santa Cruz Biotechnology (Dallas, TX); purified anti-human CD43 (1G10, #555474; RRID:AB_395866) and Alexa Fluor^®^ 647-conjugated anti-human CD3ε (UCHT1, #557706; RRID:AB_396815) from BD Sciences (San Jose, CA); Alexa Fluor^®^ 647-conjugated anti-rat Thy1 (OX-7, #202508; RRID:AB_492884) from BioLegend (San Diego, CA). Ghost Dye™ Violet 450 was from Tonbo Biosciences (#13-0863-T100; San Diego, CA). The inhibitors PP2 (#529573) and piceatannol (#527948) were from Sigma (St. Louis, MO). Linear polyethylenimine (PEI 25K, #23966) was from Polysciences, Inc (Warrington, PA).

### DNA constructs

All chimeric proteins are named with their constituent proteins listed from N-terminus to C-terminus. Expression vectors are based on pEGFP-n1 or pEGFP-c1 (Clontech, Mountain View, CA). The VSV-G pseudotyping vector pMD2.G and the viral packaging vector psPAX2 were gifts from Didier Trono (Addgene plasmids #12259 and #12260). Self-inactivating lentiviral transfer vectors are based on pLK4-EFv6-MCS-IRES-Puro, a custom vector that contains the elements of the pEGFP-n1 backbone responsible for bacterial propagation and selection and the LTR, packaging, IRES, puromycin resistance, and WPRE sequences from pLEX-MCS (Open Biosystems, Lafayette, CO). Upstream of the IRES element, pLK4-EFv6-MCS-IRES-Puro vectors incorporate a modified variant of the human EF1α enhancer and multiple cloning sites derived from either pEGFP-n1 or pEGFP-c1. Modular cassettes encoding the fluorescent proteins CFP, YFP, mYFP, and mCFP were previously inserted into these vectors. Cassettes encoding TagRFP-T (TRT) and mRFP1 were derived from material provided by R. Tsien (University of California, San Diego, La Jolla, CA) (Campbell et al., 2002; Shaner et al., 2008). The mCerulean3 (mCer3) module was derived synthetically (Markwardt et al., 2011). The tandem affinity purification tag encoding streptavidin-binding protein (SBP), a TEV protease cleavage site, and a 6xHIS tag was derived synthetically and inserted upstream of mCer3 (Keefe et al., 2001; Li et al., 2011). Vectors encoding human ZAP-70.YFP and human SLP-76.YFP have been described (Bunnell et al., 2002). The kinase dead K369R mutant of ZAP-70 was generated by site-directed mutagenesis. Fluorescent tags were exchanged using standard cloning techniques. Murine NEMO was a gift from Shao-Cong Sun (MD Anderson, Houston, TX). Human NEMO was purchased from Open Biosystems (IMAGE clone 8517). Murine Lck was provided by Lawrence Samelson (NCI, Bethesda, MD) (van Leeuwen et al., 1999). Novel chimeric inserts and NEMO truncation mutants were generated by PCR. Point mutations of murine NEMO were generated by site-directed mutagenesis. All synthetic DNAs were from IDT (Coralville, IA).

### Cell lines and primary T cell cultures

The Jurkat human leukemic T cell line E6.1 (TIB-152; RRID:CVCL_0367) and the C305 hybridoma (CRL-2424; RRID:CVCL_K130) were obtained from ATCC (Manassas, VA). Jurkat-derived lines lacking Lck (J.CaM1.6; RRID:CVCL_0354), LAT (J.CaM2.5; RRID:CVCL_DR61), and SLP-76 (J14; RRID:CVCL_R861) were provided by Arthur Weiss (UCSF, San Francisco, CA) (Finco et al., 1998; Goldsmith et al., 1988; Goldsmith and Weiss, 1987; Straus and Weiss, 1992; Yablonski et al., 1998). The ZAP-70-deficient Jurkat derivative P116 (RRID:CVCL_6429) was provided by Robert Abraham (Sanford-Burnham, La Jolla, CA) (Williams et al., 1998). The Carma1-deficient Jurkat derivative JPM50.6 was provided by Xin Lin (MD Anderson, Houston, TX) (Wang et al., 2002). NEMO-deficient Jurkat 8321 cells were provided by Jordan Orange (Baylor College of Medicine, Houston, TX) (He and Ting, 2002). The absence of all relevant proteins was confirmed by western blotting. Unless otherwise noted, Jurkat derivatives and hybridomas were maintained in complete RPMI (RPMI-1640 supplemented with 10% fetal bovine serum, 2 mM L-glutamine, and 10 μg/ml ciprofloxacin). HEK 293T cells (RRID:CVCL_0063) were obtained from Lawrence Samelson (NIH) and maintained in DMEM supplemented with 10% FBS, 2 mM L-glutamine, 100 units/ml of penicillin, and 100 μg/mL of streptomycin. De-identified human blood samples were obtained from NY Biologics Inc. (Southampton, NY). Mononuclear cells were isolated on Ficoll-Paque PLUS (GE Healthcare, Chicago, IL) and stimulated in complete RPMI using 1% phytohemagglutinin (Invitrogen). After 36-48 hr, stimulated cells were rinsed into fresh media containing 20 units/ml IL-2, and maintained in IL-2 thereafter. After 4 days, > 95% of the cells were CD3ε+, as assessed by flow cytometry.

### Production of lentiviral particles

Lentiviruses were packaged in HEK 293T cells using the VSV-G pseudotyping vector pMD2.G and the viral packaging vector psPAX2 (gifts from Didier Trono; Addgene plasmids #12259 and #12260). Transfection mixes were prepared by combining the transfer, packaging, and pseudotyping vectors with 1 mg/ml linear polyethylenimine in DMEM. These mixtures were then added dropwise to sub-confluent HEK 293T cells. The transfection medium was removed after 12-16 hr and viral supernatants were harvested 24-36 hr thereafter. Residual packaging cells were removed using 0.45 μm filters. If necessary, viral supernatants were concentrated using an Amicon centrifugal filter unit (50,000 MWCO). All procedures involving lentiviral particles were performed with the approval of the Tufts Institutional Biosafety Committee.

### Transient transfection and lentiviral transduction of T cells

Transfections and transductions were performed using mid-log phase Jurkat cells or T cell blasts 6-8 days post-PHA stimulation. For transfections, T cells were rinsed into fresh medium and briefly incubated with the desired DNA. Jurkat T cells were resuspended at 4 × 10^7^ cells/ml and T cell blasts were resuspended at 10^8^ cells/ml. Transfections were performed in 4 mm-gap cuvettes using a BTX ECM 830 square wave electroporator. Jurkat cells were transfected using a single pulse of 300 volts and 10 ms, with 5-10 μg of DNA per 100 μl of cells. T cell blasts were transfected using a single pulse of 385 volts and 6 ms, with 25 μg of DNA per 100 μl of cells (Nguyen et al., 2008). Transfected T cells were analyzed by flow cytometry and imaged within 6-48 hr. T cells were transduced by combining equal parts of mid-log phase cells, lentiviral supernatants, and complete RPMI. For the transduction of T cell blasts, the transduction mixture was supplemented with IL-2. After 2-3 days residual viruses were removed and fixed cells were analyzed by flow cytometry. If required, stably transduced T cells were enriched by selection in puromycin (1 μg/ml) or zeocin (300 μg/ml).

### T Cell Stimulation

All imaging assays were performed in 96-well glass-bottomed plates (Whatman/Matrix Technologies, Indianapolis, IN). Plates were coated with 0.01% poly-L-Lysine (Sigma) for 5 min at room temperature and then dried for 30 min at 42°C. Proteins were adsorbed onto the plate by incubating for 60 min at 37°C. First, plates were coated with 10μg/ml of OKT3. Second, the plates were co-coated with adhesive and costimulatory ligands, as required. Recombinant human VCAM-1 was used at 1μg/ml and anti-human CD28 was used at 3 μg/ml. Finally, the plates were blocked with 1% BSA to limit the deposition of serum components during imaging. Coated wells were filled with PBS and stored at 4°C for up to 3 days before use. For live cell imaging runs, pre-warmed plates were mounted on the imaging system and maintained at 37°C (Bunnell et al., 2003). Prior to each imaging run, the relevant well was rinsed, filled with HEPES-buffered complete culture medium (25 mM, pH 7.4), and allowed to equilibrate to 37°C. Imaging runs were initiated by injecting T cells into these wells. T cells typically settle onto the stimulatory substrate within 2 min and reach peak spreading within 3 min of initial contact. Unless otherwise noted, cells were imaged continuously for 5 min, beginning either at the first moment of contact or after achieving peak spreading. For fixed-cell studies, plates were equilibrated in a tissue culture incubator and cells were fixed with 1% paraformaldehyde 7-10 min after injection.

### Image acquisition

Imaging runs were performed on a modified confocal imaging system (UltraVIEW; PerkinElmer, Waltham, MA) incorporating a spinning-disc head (CSU-10; Yokogawa Electric, Japan), an Axiovert 200M stand (Carl Zeiss, Germany), and a PIFOC piezoelectric Z-drive (Physik Instrumente, Germany). Images were acquired using 40× Plan-Neofluar (NA 1.3) or 63× Plan-Apochromat (NA 1.4) oil immersion lenses, with or without a 1.6× Optovar lens (Carl Zeiss). Laser excitation was provided via a 5-line launch (442/488/514/568/647 nm; Prairie, Rochester, MN), including HeCd, Ar, and Kr/Ar gas lasers (Melles Griot, Carlsbad, CA), or via a 4-line launch based on diode and DPSS lasers (445/515/561/642 nm; Oxxius, France). Gas and DPSS laser outputs were controlled via acousto-optical modulation and diode lasers were controlled directly. CFP and YFP were imaged using a dual-pass (442/514 nm) dichroic and emission windows of 480/30 nm (CFP) and 535/30 nm (YFP). YFP was imaged with TRT, mRFP1, or FM 4-64 using a ‘RGBA’ quad-pass (405/488/568/647 nm) dichroic and emission windows of 525/50 nm (YFP), 600/50 nm (TRT or mRFP1), and 700/75 nm (FM 4-64). Cells coexpressing CFP and mRFP1 were imaged using a ‘multi-pass’ (457/514/568/647 nm) dichroic and emission windows of 480/30 nm (CFP), 535/30 nm (YFP), and 600/50 nm (mRFP1). Emission filter wheels were from Sutter (Lambda 10-2; Sacramento, CA) or ASI (FW-1000; Eugene, OR). Images were acquired using a CCD camera (Orca-II-ER; Hamamatsu Photonics, Japan), an intensified CCD camera (XR MEGA-10; Stanford Photonics, Palo Alto, CA), or a digital CMOS camera (Orca-Flash4.0 v2; Hamamatsu Photonics). The intensified CCD camera was coupled via a 2.5× expanding lens to provide resolution comparable to the Orca-II-ER. The same expanding lens was used to improve filling of the CMOS camera. System hardware was controlled using proprietary software from PerkinElmer (UltraView Imaging Suite 5.2) or using digital-analog output devices from Data Translations Inc. (DT9812; Marlborough, MA) and National Instruments (NI-6713 DAQ; Woburn, MA) in conjunction with the open-source Micro-Manager software package (Edelstein et al., 2010). For live cell studies samples were maintained at 37°C by establishing a stable temperature gradient using an air blower (Nevtek, Williamsville, VA) and a lens heater (Bioptechs, Butler, PA).

### Image processing and analysis

Individual TIFF images were combined into stacks using iVision 4.5 (BioVision Technologies, Chester Springs, PA) or the Fiji distribution of ImageJ (Schindelin et al., 2012; Schneider et al., 2012). Complex workflows were managed using AppleScript (Apple, Cupertino, CA). All subsequent image processing was performed using iVision. All images have been subjected to standard thresholding and scaling. Supplemental movies were exported using the highest quality TIFF compressor in the ‘Export Sequence to Movie’ tool. Kymographs and threedimensional projections were generated using ‘3D Projector’ tool. Line scans were generated using the ‘Extract ROI Values’ tool. In preparation for cluster identification, all images were scaled to a resolution of 100 nm/pixel.

To facilitate cluster identification across a wide range of intensities, raw images were subjected to a custom ‘Local Background’ linear filter, which emphasizes regions with sharp transitions in intensity by reducing the value of each pixel by the average value of all pixels 200 nm removed from the original. Provisional clusters were identified by manually thresholding the filtered image. Final cluster masks were generated by subjecting the provisional mask to a 3×3 median filter in order to remove noise. Static microclusters were identified in the same manner as mobile clusters, but using an ‘average-over-time’ image generated using the ‘3D Stacked View’ tool. Because the static microclusters detected in this manner are obscured if transiently occluded by brighter macroclusters, the raw image stack was pre-processed to remove macroclusters before generating the ‘average-over-time’ image. First, cluster masks larger than 0.4 μm^2^ (an effective diameter of 0.71 μm) were identified in each frame of the raw stack. Second, these masks were dilated five times to encompass the ‘halo’ that surrounds each macrocluster. Finally, the corresponding pixels inside these enlarged masks were set to the average cellular intensity outside of these masks. Data tables recording the areas, intensities, and positions of static and mobile clusters were generated using the ‘Measure Segments’ tool. To enable background corrections and the normalization of values by cellular totals, intensities and areas were also recorded for a nearby cell-free region and for a mask encompassing the relevant cell. To quantitate experimental changes in the distribution of NEMO clusters, objects were binned by area using the ‘Classify Segments’ tool. The total number of objects, the total intensity, and the total area within each class were recorded at each timepoint. For live image stacks, the resulting data were averaged over time, yielding a single data set per cell. Because mobile clusters undergo frequent changes in their direction of movement, and may merge with static microclusters or with macroclusters, we developed a user-assisted algorithm to generate NEMO cluster trajectories. This algorithm links clusters between frames using a score based on their relative positions, areas, and intensities. Trajectory formation continues automatically unless multiple pairings yield similar scores or until the score falls below an arbitrary threshold. In the former case the top matches are presented for user input. In the latter case the trajectory is terminated. Instantaneous speeds were calculated using the distance between linked clusters and the time interval between frames; peak speeds were recorded for each trajectory.

### Fluorescence Resonance Energy Transfer (FRET) Assays

FRET assays addressing the basal interactions among NEMO variants expressed in Jurkat cells were performed using an LSR II flow cytometer (BD Biosciences). Jurkat cells were transfected or transduced with pairs of mCFP- and mYFP-tagged chimeras. Fluorescent signals were monitored using the following excitation laser lines and emission windows: CFP, 405/425-475 nm; YFP 488/505-550 nm; and FRET, 405/515-545 nm. To control for variations in donor and acceptor expression within the resulting bulk populations, FRET signals were calculated for cells falling within narrowly defined CFP and YFP gates common to all samples within an experiment. This ensures that the terms contributed by CFP bleed-through and YFP cross-excitation are effectively constant, and can be eliminated by normalizing the FRET signal to the mCFP signal and subtracting the equivalent value from a non-interacting pair. In all experiments, mCFP and mYFP.NEMO were taken as the non-interacting pair.

### Cell lysis and NEMO co-precipitation

Jurkat cells were lysed at 2.5 × 10^7^ cells/ml in a buffer containing 1% Triton X-100, 20 mM Tris-HCl, pH 7.4, 100 mM NaCl, 10 mM NaF, 1 mM Na_3_VO_4_, and the EDTA-free cOmplete cocktail of protease inhibitors (Roche Applied Science, Germany). Lysates were clarified by centrifugation after 10 min on ice. YFP.NEMO chimeras were immunoprecipitated using anti-GFP (ab290) and protein A agarose beads (Thermo Fisher, Waltham, MA). Chimeras bearing tandem affinity purification tags were captured using streptavidin-sepharose beads (GE Healthcare). SDS-PAGE gels were transferred to either Immobilon-P (EMD Millipore, Billerica, MA) or nitrocellulose (Bio-Rad, Hercules, CA) membranes using a semi-dry transfer apparatus. Western blots were developed by enhanced chemiluminescence.

### NF-κB reporter assays

Jurkat 8321 cells are NEMO-deficient and stably express a reporter construct with rat Thy1 under the control of eight tandem copies of a NF-κB enhancer (He & Ting, 2002). Jurkat 8321 cells were grown in RPMI media supplemented with 10% bovine growth serum (Hyclone), 100 units/ml of penicillin, 100 μg/ml streptomycin and 1 mM L-glutamine. Mid-log phase cells were transfected by electroporation using a Bio-Rad GenePulser XCell set at 250 V and 950 μF. Cells were transfected in 4 mm-gap cuvettes with 15 μg of mYFP.NEMO expression plasmids per cuvette. The following day, transfected cells at 2.5 × 10^6^ cells/ml were plated at 200 μl/well in 96-well flat-bottom plates. Plated cells were stimulated with 20 ng/ml phorbol 12-myristate 13-acetate (PMA; Sigma) and 1 μg/ml ionomycin (Fisher Scientific, Pittsburgh, PA), 20 ng/ml PMA and 1 μg/ml anti-CD28 (9.3), or anti-TCR (C305) and 1 μg/ml anti-CD28. After 24 hr, the mean fluorescence intensity of Thy1 in YFP-gated cells was determined by flow cytometry.

### Figure Preparation and Statistical Analyses

All graphs and imaging figures were prepared with Microsoft Excel, PowerPoint, and iVision. Statistical significances were calculated using Student’s t tests for unpaired samples with equal variances. Statistical comparisons of microcluster populations are presented in detail in Supplemental Table 1, which also shows effect sizes (Cohen’s d).

## Supplemental Materials

One Supplemental Table, five Supplemental Figures and twenty-two Supplemental Movies are associated with this manuscript.

## Acknowledgements

We thank Warner Greene and Shao Cong-Sun for the IKKγ cDNA; Shao Cong-Sun for NEMO-deficient Jurkat T cells; Xin Lin for the Carma1-deficient Jurkat T cells. We also thank the Tufts University Flow Cytometry Core. This work was supported by NIH grant GM080398 (to L.P.K.), an ARRA supplement to GM080398 (to L.P.K. and S.C.B.) and NIH grant AI103022 (to L.P.K. and S.C.B.). In addition, we recognize the W.M. Keck Foundation and the Eshe Fund for their generous support of core facilities at Tufts.

## Author Contributions

Conceptualization: LPK and SCB; Methodology: LPK and SCB; Software, SCB; Formal Analysis: EAD, ALS, AM, and SCB; Investigation: EAD, ALS, AM, JMM, MS, LPK and SCB; Writing – Original Draft: EAD; Writing – Review & Editing: ALS, MS, LPK, and SCB; Visualization: EAD and SCB; Supervision, Project Administration, and Funding Acquisition: LPK and SCB.

## Supplemental Figure Legends

**Supplements to Figure 1:**

A) Jurkat T cells were stably transduced with wild-type murine or human mYFP-NEMO chimeras and sorted to obtain levels of mYFP fluorescence. Comparable levels of chimera expression were confirmed by western blotting (n=3 experiments). B) Both human and murine NEMO co-immunoprecipitate endogenous human IKKβ from Jurkat cells stably transduced with NEMO.mYFP chimeras (n=3 experiments). C) A transiently transfected murine NEMO chimera reconstitutes NF-κB dependent signals in Jurkat derived, NEMO-deficient 8321 cells bearing an NF-κB-driven rat Thy1 reporter. These cells were left unstimulated (n=5) or stimulated with PMA and ionomycin (P+I; n=5), PMA and anti-CD28 (P+28; n=4), or anti-CD3 and anti-CD28 (TCR+28; n=2). Data are presented as the mean ± SEM. Statistical differences were determined using Student’s T test: p < 0.01, **; p < 0.001, ***. D) Jurkat T cells expressing murine (left panel) or human (right panel) mYFP.NEMO chimeras were stimulated on substrates coated with anti-CD3 and rhVCAM1, as in Fig. 1 (n=3 experiments). E) Jurkat T cells stably expressing murine mYFP.NEMO were stimulated on fibronectin or anti-CD3, as in Figure 1A. All images and projections are derived from confocal Z-stacks acquired at peak spreading (n=2 experiments). F) Representative kymographs used for the derivation of the lag time to NEMO microcluster formation in Figure 1C. Red brackets span the period from contact initiation to microcluster nucleation. Red arrows identify mobile NEMO structures. Scale bars: 10 μm (stills), 5 μm × 60 seconds (kymographs), 10 μm × 10 μm (Z-projections).

**Supplements to Figure 3:**

A) FRET-based analysis of NEMO oligomerization in live cells. Jurkat T cells were stably transduced or transiently transfected with the indicated mCFP-tagged donor and mYFP-tagged acceptor constructs and analyzed for FRET (see Materials and Methods). Background-corrected FRET signals are expressed as a percentage of the response observed with a wild-type donor and acceptor. Data are presented as the mean ± SEM, based on the number of independent experiments. Differences from WT were assessed using Student’s T-test. A graphic summary of the results is presented at right. B) Biochemical analysis of NEMO oligomerization in cell lysates. 293T cells were stably transduced with the indicated NEMO constructs, lysed, and captured using streptavidin agarose. Data are representative of at least three experiments.

**Supplements to Figure 4:**

A) Jurkat T cells were stably transduced or transiently transfected with a mCFP.NEMO donor construct and the indicated mYFP-tagged acceptor constructs and analyzed for FRET as in Supplemental Figure 4A. Data are presented as the mean ± SEM, based on the number of independent experiments. Differences from WT were assessed using Student’s T-test. B) Biochemical analysis of NEMO oligomerization in cell lysates. 293T cells were stably transduced with the indicated NEMO constructs, lysed, and captured using streptavidin agarose. Data are representative of at least three experiments. C) Co-immunoprecipitation of endogenous human IKKβ with mYFP.NEMO chimeras exogenously expressed in Jurkat T cells. Data are representative of at least three experiments. D) Relative fluorescence intensity profiles (along white diagonal lines) are shown for the primary T cells imaged in Figure 5E. Scale bars: 10 μm.

## Supplemental Movie Legends

**Supplemental Movies 1-3**

Jurkat T cells were transiently transfected with vectors encoding mYFP.NEMO, stimulated on the indicated substrates, as in Figure 1A, and imaged continuously for 5 minutes. Movie 1: Fibronectin; Movie 2: Anti-CD3; Movie 3: Anti-CD3 and rhVCAM1. Scale bars: 10 μm. Compression Rate: 60x.

**Supplemental Movie 4**

Primary human T cell blasts were transiently transfected with vectors encoding mYFP.NEMO, stimulated on anti-CD3, anti-CD28, and rhVCAM1, as in Figure 1B, and imaged continuously for 5 minutes. Scale bars: 10 μm. Compression Rate: 60x.

**Supplemental Movie 5**

Jurkat T cells were transiently transfected ZAP-70.CFP (red) and mYFP.NEMO (green), stimulated on anti-CD3, and imaged for 5 minutes. The same cell is shown in Figure 2B. Scale bars: 10 μm. Compression Rate: 60x.

**Supplemental Movies 6-9**

Jurkat T cells stably transduced with variants of mYFP.NEMO were stimulated on anti-CD3 and rhVCAM1 and imaged for 5 minutes. The same cells are shown in Figure 5C. Movie 13: wild-type NEMO; Movie 14: NEMO Y301S mutant; Movie 15: NEMO R312Q mutant; Movie 16: NEMO V407S mutant. Scale bars: 10 μm. Compression Rate: 60x.

**Supplemental Movies 10-11** Jurkat T cells stably transduced with variants of mYFP.NEMO were pre-incubated with 10 μM PP2 or a corresponding volume of DMSO for 10 minutes and stimulated on anti-CD3 and rhVCAM1 in the presence of the corresponding compound. Cells were imaged for 5 minutes, beginning within 2 minutes of cell addition. The movies correspond to the kymographs shown in the middle panel of Figure 6B. Movie 17: DMSO treated; Movie 18: PP2 treated. Scale bars: 10 μm. Compression Rate: 60x.

**Supplemental Movies 12-13**

ZAP-70-deficient Jurkat P116 cells were transiently transfected with vectors encoding mYFP.NEMO and either a wild-type (WT) or a kinase-dead (K369R, KR) ZAP-70.mCFP chimera. Cells were stimulated on substrates coated with anti-CD3, rhVCAM1, and anti-CD28 and imaged continuously for 5 minutes. The same cells are shown in Figure 6D. Movie 19: NEMO with wild-type ZAP-70; Movie 20: NEMO with kinase-dead ZAP-70. Scale bars: 10 μm. Compression Rate: 60x.

## References

Bagneris, C., Ageichik, A.V., Cronin, N., Wallace, B., Collins, M., Boshoff, C., Waksman, G., and Barrett, T. (2008). Crystal structure of a vFlip-IKKγ complex: insights into viral activation of the IKK signalosome. Mol Cell 30, 620–631.

Bunnell, S.C. (2010). Multiple microclusters: diverse compartments within the immune synapse. Current topics in microbiology and immunology 340, 123–154.

Bunnell, S.C., Barr, V.A., Fuller, C.L., and Samelson, L.E. (2003). High-resolution multicolor imaging of dynamic signaling complexes in T cells stimulated by planar substrates. Science’s STKE: signal transduction knowledge environment 2003, PL8.

Bunnell, S.C., Hong, D.I., Kardon, J.R., Yamazaki, T., McGlade, C.J., Barr, V.A., and Samelson, L.E. (2002). T cell receptor ligation induces the formation of dynamically regulated signaling assemblies. J Cell Biol 158, 1263–1275.

Campbell, R.E., Tour, O., Palmer, A.E., Steinbach, P.A., Baird, G.S., Zacharias, D.A., and Tsien, R.Y. (2002). A monomeric red fluorescent protein. Proc Natl Acad Sci U S A 99, 7877–7882.

Campi, G., Varma, R., and Dustin, M.L. (2005). Actin and agonist MHC-peptide complex-dependent T cell receptor microclusters as scaffolds for signaling. J Exp Med 202, 1031–1036.

Cheng, J., Montecalvo, A., and Kane, L.P. (2011). Regulation of NF-κB induction by TCR/CD28. Immunol Res 50, 113–117.

Coornaert, B., Baens, M., Heyninck, K., Bekaert, T., Haegman, M., Staal, J., Sun, L., Chen, Z.J., Marynen, P., and Beyaert, R. (2008). T cell antigen receptor stimulation induces MALT1 paracaspase-mediated cleavage of the NF-κB inhibitor A20. Nat Immunol 9, 263–271.

Cordier, F., Grubisha, O., Traincard, F., Veron, M., Delepierre, M., and Agou, F. (2009). The zinc finger of NEMO is a functional ubiquitin-binding domain. J Biol Chem 284, 2902–2907.

Deng, L., Wang, C., Spencer, E., Yang, L., Braun, A., You, J., Slaughter, C., Pickart, C., and Chen, Z.J. (2000). Activation of the IkappaB kinase complex by TRAF6 requires a dimeric ubiquitin-conjugating enzyme complex and a unique polyubiquitin chain. Cell 103, 351–361.

Dienz, O., Moller, A., Strecker, A., Stephan, N., Krammer, P.H., Droge, W., and Schmitz, M.L. (2003). Src Homology 2 Domain-Containing Leukocyte Phosphoprotein of 76 kDa and Phospholipase Cγ1 Are Required for NF-κB Activation and Lipid Raft Recruitment of Protein Kinase Cθ Induced by T Cell Costimulation. J Immunol 170, 365–372.

Drew, D., Shimada, E., Huynh, K., Bergqvist, S., Talwar, R., Karin, M., and Ghosh, G. (2007). Inhibitor κB kinase β binding by inhibitor κB kinase γ. Biochemistry 46, 12482–12490.

Ea, C.K., Deng, L., Xia, Z.P., Pineda, G., and Chen, Z.J. (2006). Activation of IKK by TNFα requires site-specific ubiquitination of RIP1 and polyubiquitin binding by NEMO. Mol Cell 22, 245–257.

Edelstein, A., Amodaj, N., Hoover, K., Vale, R., and Stuurman, N. (2010). Computer control of microscopes using microManager. Curr Protoc Mol Biol Chapter 14, Unit14 20.

Filipe-Santos, O., Bustamante, J., Haverkamp, M.H., Vinolo, E., Ku, C.L., Puel, A., Frucht, D.M., Christel, K., von Bernuth, H., Jouanguy, E., et al. (2006). X-linked susceptibility to mycobacteria is caused by mutations in NEMO impairing CD40-dependent IL-12 production. J Exp Med 203, 1745–1759.

Finco, T.S., Kadlecek, T., Zhang, W., Samelson, L.E., and Weiss, A. (1998). LAT is required for TCR-mediated activation of PLCγ1 and the Ras pathway. Immunity 9, 617–626.

Fujita, H., Rahighi, S., Akita, M., Kato, R., Sasaki, Y., Wakatsuki, S., and Iwai, K. (2014). Mechanism underlying IκB kinase activation mediated by the linear ubiquitin chain assembly complex. Mol Cell Biol 34, 1322–1335.

Goldsmith, M.A., Dazin, P.F., and Weiss, A. (1988). At least two non-antigen-binding molecules are required for signal transduction by the T-cell antigen receptor. Proc Natl Acad Sci U S A 85, 8613–8617.

Goldsmith, M.A., and Weiss, A. (1987). Isolation and characterization of a T-lymphocyte somatic mutant with altered signal transduction by the antigen receptor. Proc Natl Acad Sci U S A 84, 6879–6883.

Grubisha, O., Kaminska, M., Duquerroy, S., Fontan, E., Cordier, F., Haouz, A., Raynal, B., Chiaravalli, J., Delepierre, M., Israel, A., et al. (2010). DARPin-assisted crystallography of the CC2-LZ domain of NEMO reveals a coupling between dimerization and ubiquitin binding. J Mol Biol 395, 89–104.

Hadian, K., Griesbach, R.A., Dornauer, S., Wanger, T.M., Nagel, D., Metlitzky, M., Beisker, W., Schmidt-Supprian, M., and Krappmann, D. (2011). NF-κB essential modulator (NEMO) interaction with linear and lys-63 ubiquitin chains contributes to NF-κB activation. J Biol Chem 286, 26107–26117.

Hanke, J.H., Gardner, J.P., Dow, R.L., Changelian, P.S., Brissette, W.H., Weringer, E.J., Pollok, B.A., and Connelly, P.A. (1996). Discovery of a novel, potent, and Src family-selective tyrosine kinase inhibitor. Study of Lck- and FynT-dependent T cell activation. J Biol Chem 271, 695–701.

Hanson, E.P., Monaco-Shawver, L., Solt, L.A., Madge, L.A., Banerjee, P.P., May, M.J., and Orange, J.S. (2008). Hypomorphic nuclear factor-κB essential modulator mutation database and reconstitution system identifies phenotypic and immunologic diversity. J Allergy Clin Immunol 122, 1169–1177 e1116.

Hara, H., Bakal, C., Wada, T., Bouchard, D., Rottapel, R., Saito, T., and Penninger, J.M. (2004). The molecular adapter Carma1 controls entry of IκB kinase into the central immune synapse. J Exp Med 200, 1167–1177.

Hashimoto-Tane, A., and Saito, T. (2016). Dynamic Regulation of TCR-Microclusters and the Microsynapse for T Cell Activation. Frontiers in immunology 7, 255.

Hayden, M.S., and Ghosh, S. (2012). NF-κB, the first quarter-century: remarkable progress and outstanding questions. Genes Dev 26, 203–234.

Hayden, M.S., West, A.P., and Ghosh, S. (2006). NF-κB and the immune response. Oncogene 25, 6758–6780.

He, K.L., and Ting, A.T. (2002). A20 Inhibits Tumor Necrosis Factor (TNF) Alpha-Induced Apoptosis by Disrupting Recruitment of TRADD and RIP to the TNF Receptor 1 Complex in Jurkat T Cells. Molecular and Cellular Biology 22, 6034–6045.

Herndon, T.M., Shan, X.C., Tsokos, G.C., and Wange, R.L. (2001). ZAP-70 and SLP-76 Regulate Protein Kinase C-θ and NF-κB Activation in Response to Engagement of CD3 and CD28. The Journal of Immunology 166, 5654–5664.

Israel, A. (2010). The IKK complex, a central regulator of NF-κB activation. Cold Spring Harb Perspect Biol 2, a000158.

Keefe, A.D., Wilson, D.S., Seelig, B., and Szostak, J.W. (2001). One-step purification of recombinant proteins using a nanomolar-affinity streptavidin-binding peptide, the SBP-Tag. Protein Expr Purif 23, 440–446.

Kensche, T., Tokunaga, F., Ikeda, F., Goto, E., Iwai, K., and Dikic, I. (2012). Analysis of nuclear factor-κB (NF-κB) essential modulator (NEMO) binding to linear and lysine-linked ubiquitin chains and its role in the activation of NF-κB. J Biol Chem 287, 23626–23634.

Laplantine, E., Fontan, E., Chiaravalli, J., Lopez, T., Lakisic, G., Veron, M., Agou, F., and Israel, A. (2009). NEMO specifically recognizes K63-linked poly-ubiquitin chains through a new bipartite ubiquitin-binding domain. EMBO J 28, 2885–2895.

Lee, K.H., Holdorf, A.D., Dustin, M.L., Chan, A.C., Allen, P.M., and Shaw, A.S. (2002). T cell receptor signaling precedes immunological synapse formation. Science 295, 1539–1542.

Li, Y., Franklin, S., Zhang, M.J., and Vondriska, T.M. (2011). Highly efficient purification of protein complexes from mammalian cells using a novel streptavidin-binding peptide and hexahistidine tandem tag system: application to Bruton’s tyrosine kinase. Protein Sci 20, 140–149.

Lo, Y.C., Lin, S.C., Rospigliosi, C.C., Conze, D.B., Wu, C.J., Ashwell, J.D., Eliezer, D., and Wu, H. (2009). Structural basis for recognition of diubiquitins by NEMO. Mol Cell 33, 602–615.

Markwardt, M.L., Kremers, G.J., Kraft, C.A., Ray, K., Cranfill, P.J., Wilson, K.A., Day, R.N., Wachter, R.M., Davidson, M.W., and Rizzo, M.A. (2011). An improved cerulean fluorescent protein with enhanced brightness and reduced reversible photoswitching. PLoS One 6, e17896.

Napetschnig, J., and Wu, H. (2013a). Molecular basis of NF-kappaB signaling. Annu Rev Biophys 42, 443–468.

Napetschnig, J., and Wu, H. (2013b). Molecular basis of NF-κB signaling. Annu Rev Biophys 42, 443–468.

Ngadjeua, F., Chiaravalli, J., Traincard, F., Raynal, B., Fontan, E., and Agou, F. (2013). Twosided ubiquitin binding of NF-κB essential modulator (NEMO) zinc finger unveiled by a mutation associated with anhidrotic ectodermal dysplasia with immunodeficiency syndrome. J Biol Chem 288, 33722–33737.

Nguyen, K., Sylvain, N.R., and Bunnell, S.C. (2008). T cell costimulation via the integrin VLA-4 inhibits the actin-dependent centralization of signaling microclusters containing the adaptor SLP-76. Immunity 28, 810–821.

Orange, J.S., Jain, A., Ballas, Z.K., Schneider, L.C., Geha, R.S., and Bonilla, F.A. (2004). The presentation and natural history of immunodeficiency caused by nuclear factor κB essential modulator mutation. J Allergy Clin Immunol 113, 725–733.

Palkowitsch, L., Leidner, J., Ghosh, S., and Marienfeld, R.B. (2008). Phosphorylation of serine 68 in the IkappaB kinase (IKK)-binding domain of NEMO interferes with the structure of the IKK complex and tumor necrosis factor-alpha-induced NF-kappaB activity. J Biol Chem 283, 76–86.

Paul, S., Kashyap, A.K., Jia, W., He, Y.W., and Schaefer, B.C. (2012). Selective autophagy of the adaptor protein Bcl10 modulates T cell receptor activation of NF-κB. Immunity 36, 947–958.

Paul, S., and Schaefer, B.C. (2013). A new look at T cell receptor signaling to nuclear factor-κB. Trends Immunol 34, 269–281.

Paul, S., Traver, M.K., Kashyap, A.K., Washington, M.A., Latoche, J.R., and Schaefer, B.C. (2014). T cell receptor signals to NF-κB are transmitted by a cytosolic p62-Bcl10-Malt1-IKK signalosome. Sci Signal 7, ra45.

Prajapati, S., and Gaynor, R.B. (2002). Regulation of Ikappa B kinase (IKK)gamma /NEMO function by IKKbeta-mediated phosphorylation. J Biol Chem 277, 24331–24339.

Rahighi, S., Ikeda, F., Kawasaki, M., Akutsu, M., Suzuki, N., Kato, R., Kensche, T., Uejima, T., Bloor, S., Komander, D., et al. (2009). Specific recognition of linear ubiquitin chains by NEMO is important for NF-κB activation. Cell 136, 1098–1109.

Rebeaud, F., Hailfinger, S., Posevitz-Fejfar, A., Tapernoux, M., Moser, R., Rueda, D., Gaide, O., Guzzardi, M., Iancu, E.M., Rufer, N., et al. (2008). The proteolytic activity of the paracaspase MALT1 is key in T cell activation. Nat Immunol 9, 272–281.

Rossman, J.S., Stoicheva, N.G., Langel, F.D., Patterson, G.H., Lippincott-Schwartz, J., and Schaefer, B.C. (2006). POLKADOTS are foci of functional interactions in T-cell receptor-mediated signaling to NF-κB. Mol Biol Cell 17, 2166–2176.

Ruland, J., Duncan, G.S., Wakeham, A., and Mak, T.W. (2003). Differential requirement for Malt1 in T and B cell antigen receptor signaling. Immunity 19, 749–758.

Rushe, M., Silvian, L., Bixler, S., Chen, L.L., Cheung, A., Bowes, S., Cuervo, H., Berkowitz, S., Zheng, T., Guckian, K., et al. (2008). Structure of a NEMO/IKK-associating domain reveals architecture of the interaction site. Structure 16, 798–808.

Schaefer, B.C., Kappler, J.W., Kupfer, A., and Marrack, P. (2004). Complex and dynamic redistribution of NF-κB signaling intermediates in response to T cell receptor stimulation. Proc Natl Acad Sci U S A 101, 1004–1009.

Schindelin, J., Arganda-Carreras, I., Frise, E., Kaynig, V., Longair, M., Pietzsch, T., Preibisch, S., Rueden, C., Saalfeld, S., Schmid, B., et al. (2012). Fiji: an open-source platform for biological-image analysis. Nat Methods 9, 676–682.

Schneider, C.A., Rasband, W.S., and Eliceiri, K.W. (2012). NIH Image to ImageJ: 25 years of image analysis. Nat Methods 9, 671–675.

Senegas, A., Gautheron, J., Maurin, A.G., and Courtois, G. (2015). IKK-related genetic diseases: probing NF-κB functions in humans and other matters. Cell Mol Life Sci 72, 1275–1287.

Shambharkar, P.B., Blonska, M., Pappu, B.P., Li, H., You, Y., Sakurai, H., Darnay, B.G., Hara, H., Penninger, J., and Lin, X. (2007). Phosphorylation and ubiquitination of the IκB kinase complex by two distinct signaling pathways. EMBO J 26, 1794–1805.

Shaner, N.C., Lin, M.Z., McKeown, M.R., Steinbach, P.A., Hazelwood, K.L., Davidson, M.W., and Tsien, R.Y. (2008). Improving the photostability of bright monomeric orange and red fluorescent proteins. Nat Methods 5, 545–551.

Shi, J.H., and Sun, S.C. (2015). TCR signaling to NF-κB and mTORC1: Expanding roles of the CARMA1 complex. Mol Immunol 68, 546–557.

Solt, L.A., Madge, L.A., and May, M.J. (2009). NEMO-binding domains of both IKKα and IKKβ regulate IκB kinase complex assembly and classical NF-κB activation. J Biol Chem 284, 27596–27608.

Solt, L.A., Madge, L.A., Orange, J.S., and May, M.J. (2007). Interleukin-1-induced NF-κB activation is NEMO-dependent but does not require IKKβ. J Biol Chem 282, 8724–8733.

Srivastava, R., Burbach, B.J., and Shimizu, Y. (2010). NF-kB Activation in T Cells Requires Discrete Control of IkB Kinase α/β (IKKα/β) Phosphorylation and IKKγ Ubiquitination by the ADAP Adapter Protein. Journal of Biological Chemistry 285, 11100–11105.

Straus, D.B., and Weiss, A. (1992). Genetic evidence for the involvement of the lck tyrosine kinase in signal transduction through the T cell antigen receptor. Cell 70, 585–593.

Sun, L., Deng, L., Ea, C.K., Xia, Z.P., and Chen, Z.J. (2004). The TRAF6 ubiquitin ligase and TAK1 kinase mediate IKK activation by BCL10 and MALT1 in T lymphocytes. Mol Cell 14, 289–301.

Tarantino, N., Tinevez, J.Y., Crowell, E.F., Boisson, B., Henriques, R., Mhlanga, M., Agou, F., Israel, A., and Laplantine, E. (2014). TNF and IL-1 exhibit distinct ubiquitin requirements for inducing NEMO-IKK supramolecular structures. J Cell Biol 204, 231–245.

Thome, M., Charton, J.E., Pelzer, C., and Hailfinger, S. (2010). Antigen receptor signaling to NF-κB via CARMA1, BCL10, and MALT1. Cold Spring Harb Perspect Biol 2, a003004.

Tokunaga, F., Sakata, S., Saeki, Y., Satomi, Y., Kirisako, T., Kamei, K., Nakagawa, T., Kato, M., Murata, S., Yamaoka, S., et al. (2009). Involvement of linear polyubiquitylation of NEMO in NF-κB activation. Nat Cell Biol 11, 123–132.

van Leeuwen, J.E., Paik, P.K., and Samelson, L.E. (1999). The oncogenic 70Z Cbl mutation blocks the phosphotyrosine binding domain-dependent negative regulation of ZAP-70 by c-Cbl in Jurkat T cells. Mol Cell Biol 19, 6652–6664.

Wang, D., You, Y., Case, S.M., McAllister-Lucas, L.M., Wang, L., DiStefano, P.S., Nunez, G., Bertin, J., and Lin, X. (2002). A requirement for CARMA1 in TCR-induced NF-κB activation. Nat Immunol 3, 830–835.

Weil, R., Schwamborn, K., Alcover, A., Bessia, C., Di Bartolo, V., and Israel, A. (2003). Induction of the NF-κB cascade by recruitment of the scaffold molecule NEMO to the T cell receptor. Immunity 18, 13–26.

Williams, B.L., Schreiber, K.L., Zhang, W., Wange, R.L., Samelson, L.E., Leibson, P.J., and Abraham, R.T. (1998). Genetic evidence for differential coupling of Syk family kinases to the T-cell receptor: reconstitution studies in a ZAP-70-deficient Jurkat T-cell line. Mol Cell Biol 18, 1388–1399.

Wu, C.J., Conze, D.B., Li, T., Srinivasula, S.M., and Ashwell, J.D. (2006). Sensing of Lys 63-linked polyubiquitination by NEMO is a key event in NF-κB activation [corrected]. Nat Cell Biol 8, 398–406.

Yablonski, D., Kuhne, M.R., Kadlecek, T., and Weiss, A. (1998). Uncoupling of nonreceptor tyrosine kinases from PLC-γ1 in an SLP-76-deficient T cell. Science 281, 413–416.

Yokosuka, T., Kobayashi, W., Sakata-Sogawa, K., Takamatsu, M., Hashimoto-Tane, A., Dustin, M.L., Tokunaga, M., and Saito, T. (2008). Spatiotemporal regulation of T cell costimulation by TCR-CD28 microclusters and protein kinase Cθ translocation. Immunity 29, 589–601.

Yokosuka, T., Sakata-Sogawa, K., Kobayashi, W., Hiroshima, M., Hashimoto-Tane, A., Tokunaga, M., Dustin, M.L., and Saito, T. (2005). Newly generated T cell receptor microclusters initiate and sustain T cell activation by recruitment of Zap70 and SLP-76. Nat Immunol 6, 1253–1262.

Yoshikawa, A., Sato, Y., Yamashita, M., Mimura, H., Yamagata, A., and Fukai, S. (2009). Crystal structure of the NEMO ubiquitin-binding domain in complex with Lys 63-linked diubiquitin. FEBS Lett 583, 3317–3322.

Zhou, H., Wertz, I., O’Rourke, K., Ultsch, M., Seshagiri, S., Eby, M., Xiao, W., and Dixit, V.M. (2004). Bcl10 activates the NF-κB pathway through ubiquitination of NEMO. Nature 427, 167–171.

